# Glucosylation of Endogenous Haustorium-inducing factors Underpins Kin Avoidance in Parasitic Plants

**DOI:** 10.1101/2025.04.17.649457

**Authors:** Lei Xiang, Songkui Cui, Simon B. Saucet, Moe Takahashi, Shoko Inaba, Bing Xie, Mario Schilder, Shota Shimada, Mengqi Cui, Mutsumi Watanabe, Yuki Tobimatsu, Harro J. Bouwmeester, Takayuki Tohge, Ken Shirasu, Satoko Yoshida

**Affiliations:** Nara Institute of Science and Technology, Graduate School of Science and Technology; Ikoma, Nara, 630-0192 Japan; Department of Economic Plants and Biotechnology, Yunnan Key Laboratory for Wild Plant Resources, Kunming Institute of Botany, Chinese Academy of Sciences; Kunming, China; RIKEN Center for Sustainable Resource Science; Yokohama, Kanagawa, Japan; Research Institute for Sustainable Humanosphere, Kyoto University, Gokasho, Uji, Kyoto 611-0011, Japan; Swammerdam Institute for Life Sciences, University of Amsterdam, Science Park 904, 1098 XH Amsterdam, the Netherlands

## Abstract

Parasitic plants rarely attack themselves, suggesting the activity of a kin-avoidance mechanism. In the root parasitic plant *Phtheirospermum japonicum*, prehaustorium formation is triggered by host-secreted haustorium-inducing factors (HIFs), but it is unresponsive to its own root exudates. Here we report the identification of the *<u>spo</u>ntaneous pre<u>h</u>austorium 1* (*spoh1*) mutant, which forms prehaustoria without external HIFs. *spoh1* harbors a point mutation in the gene encoding UDP-glycosyltransferase GT72B1, an enzyme that glucosylates and thereby inactivates phenolic HIFs. Notably, PjGT72B1 possesses a different substrate specificity from its ortholog of the host Arabidopsis. Introduction of PjGT72B1 in Arabidopsis reduced HIF activity, indicating that HIF glucosylation regulates haustorium induction by hosts. Our findings suggest that parasitic plants have evolved kin-avoidance mechanisms through the glucosylation of endogenous HIFs.

## Main Text

Parasitic plants are recognized as agricultural weeds that parasitize crop species and result in more than 1 billion USD annual economic losses, threatening global food security (*1, 2*).

Parasitic plants invade hosts and obtain nutrients through a specialized organ known as a haustorium (*3*). This multifunctional organ facilitates attachment, penetration, and connection to the host plant’s vascular system, enabling the transfer of water and nutrients. While the host range of parasitic plants vary depending on the species, previous studies have shown that parasitic plants limit the formation of this intimate connection with other individuals of the same species or other parasites in the same family, a phenomenon referred to as kin avoidance (or avoidance of self-parasitism) (*4, 5*). However, the molecular mechanisms underlying kin avoidance remain elusive.

The initiation of haustorium formation is a crucial early step in parasitism, triggered by host-derived chemical cues known as haustorium-inducing factors (HIFs). The detection of HIFs stimulates the development of prehaustoria, which are preliminary haustorial structures that eventually mature into fully functional haustoria (*3, 6*–*9*). Quinones and phenolics, such as 2,6-dimethoxy-1,4-benzoquinone (DMBQ), syringic acid, vanillin and ferulic acid (FerA) are known to act as HIFs for Orobanchaceae hemiparasitic plants (*8, 10*). These molecules are biosynthetic precursors or metabolic derivatives of cell wall lignin, a structural polymer ubiquitous among vascular plants (*10*). Lignin is produced by polymerization of monolignols, which are biosynthesized through the phenylpropanoid pathway. Depending on the number of methoxy groups at the 3- and 5-positions of the phenolic ring, lignin polymer units and lignin-associated metabolites are categorized as p-hydroxyphenyl (H)-, guaiacyl (G)- or syringyl (S)-types, containing 0, 1 and 2 methoxy groups, respectively. Notably, G- and S-type lignin precursors or related phenolics act as HIFs, with S-type compounds generally exhibiting stronger activity than G-type ones (fig S1) (*10*).

Given the ubiquitous presence of lignin compounds in vascular plants, it is likely that HIF-related compounds are also present in parasitic plant species. Intriguingly, however, Orobanchaceae parasitic plants rarely initiate prehaustorium formation in response to signals from other parasitic members of the same family (*4, 5, 11*). For example, the obligate hemiparasite *Striga hermonthica* (purpule witchweed) does not form prehaustoria when exposed to the facultative parasitic plant *Phtheirospermum japonicum* (*12*). Similarly, the facultative parasitic plants *Triphysaria* spp. rarely form self- or conspecific prehaustoria (*4, 13*).

Nevertheless, treatment with phenol oxidase enhances the release of HIFs from *Triphysaria versicolor* roots, suggesting that parasitic plants can produce HIFs but either their release or activity is suppressed by unknown mechanisms (*13*).

### Isolation of the *P. japonicum* spontaneous prehaustoria formation (*spoh1*) mutant

To investigate kin-avoidance mechanisms, we isolated mutants that form prehaustoria in the absence of external HIFs. Wild-type (WT) *P. japonicum* seedlings exhibit straight radicle growth in water incubation without HIFs, but exposure to exogenous DMBQ results in shortened, curved radicles with bump-like prehaustorial structures (fig S2). Through forward genetic screening of ethyl methyl sulfonate (EMS)-induced mutant pools of *P. japonicum* (*14*), we identified a mutant line, designated as *spontaneous prehaustorium 1* (*spoh1*), that phenocopies DMBQ-treated WT in water (fig S2). Because sucrose enhances spontaneous prehaustorium formation in *Pedicularis kansuensis*, an Orobanchaceae parasite (*15*), we used sucrose-containing agar medium for our assays. WT plants failed to produce prehaustoria after a two-week incubation on media containing 4% sucrose without external HIFs, but *spoh1* seedlings consistently generated prehaustorium-like structures (Fig. 1A). The internal structures of *spoh1* prehaustoria was indistinguishable from those of DMBQ-induced WT prehaustoria, both exhibiting a semi-spherical structure covered with densely curved haustorial hairs (Fig. 1B). Moreover, when *spoh1* plants infected Arabidopsis, the mature haustoria were morphologically indistinguishable from those of WT, indicating that the *spoh1* mutation does not affect haustorial maturation (Fig. 1C).

**Fig. 1.**
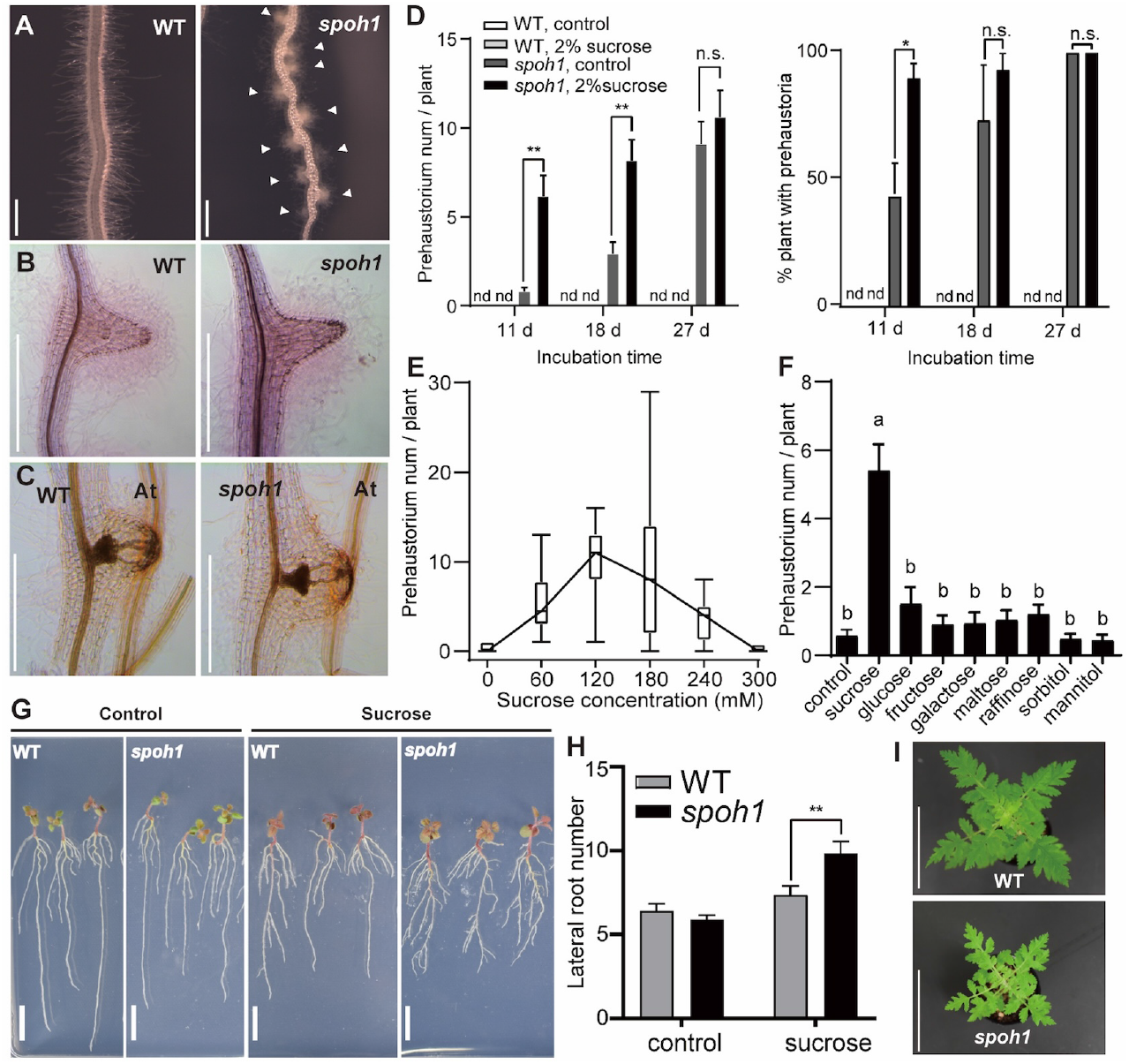
*P. japonicum spoh1* forms spontaneous prehaustoria enhanced by exogenous sucrose. (**A**) Root phenotype of WT and *spoh1* after two-week incubation on 4% sucrose-containing agar medium. White arrowheads indicate spontaneous prehaustoria in *spoh1*. (**B**) Safranin-O stain of prehaustoria of WT induced by a HIF and of *spoh1* formed spontaneously. (**C**) Safranin O stain of WT and *spoh1* mature haustoria. Scale bar: 500 μm (A-C). **(D)** Prehaustorium number per plant (left) and percentage of prehaustorium-forming plants (right) in WT and *spoh1* grown in water agar (control) or with 2% sucrose media. Data represent mean ± SE from three representative experiments; n = 30. Student’s *t* test, unpaired, two-tailed (*p < 0.05). (**E**) *spoh1* prehaustorium number after incubation for 2 weeks on different concentration of sucrose media. Data are shown as a box plot. (**F**) Prehaustoria number per plant in *spoh1*incubated on media containing 2% of each sugar for 2 weeks. Data represent mean ± SE from three representative experiments; n = 30. Different alphabets represent significant difference detected by one-way ANOVA (p<0.05). (**G**) WT and *spoh1* phenotype after incubation on media with or without 2% sucrose for 2 weeks. Scale bar: 1 cm. **(H)** Lateral root number per plant in WT and spoh1 grown as (G). Student’s t test, unpaired, two-tailed (\*\**p* < 0.01, n=30). **(I)** Crowns of 45-day WT and *spoh1* grown on soil. Scale bar: 5 cm.

While not essential for spontaneous prehaustorium formation in *spoh1*, sucrose acts as a potent enhancer. In the absence of sucrose, all *spoh1* seedlings formed about the same number of spontaneous prehaustoria within 27 days after germination. WT did not produce prehaustoria, regardless of sugar concentration, but supplementing the medium with 2% sucrose accelerated prehaustorium formation in *spoh1* (Fig. 1D). The optimal enhancement was achieved at 120 mM sucrose (approximately 4%), which yielded the highest prehaustorium number per plant (Fig. 1E). Importantly, no other sugars nor sugar alcohols replicated this effect (Fig. 1F), indicating that the promotional effect of sucrose is due to its function as a signaling molecule rather than a nutritional or osmotic pressure (*16*). Under 2% sucrose conditions, *spoh1* plants showed a modest yet significant reduction in primary root length alongside an increase in lateral root formation, compared to WT (Fig. 1G, H, fig. S3A, B). Furthermore, when grown in pots, *spoh1* displayed reduced crown diameters and shorter overall heights at 45- and 75-days, suggestive of a slight growth repression compared to WT (Fig. 1I, fig. S3C,D).

### *spoh1* root exudates induce *S. hermonthica* prehaustorium formation

To test if *spoh1* plants autonomously produce HIFs that self-induce prehaustoria, root extracts or exudates from *spoh1* and WT seedlings were applied to WT *P. japonicum* and to *S. hermonthica*, a parasitic species in the same Orobanchaceae family. Little prehaustoria-inducing activity was observed after application of root extracts from either WT nor *spoh1* seedlings incubated on media with sucrose, whereas host Arabidopsis root extracts induced approximately 15 prehaustoria per WT *P. japonicum* plant (Fig 2A). Similarly, less than 20% of *Striga* seedlings formed prehaustoria in response to WT or *spoh1* root extracts, although *spoh1* prehaustorium formation rates were higher than WT under sucrose-free conditions (Fig 2B). These results suggest that *spoh1* root extracts contain low levels of HIFs. Remarkably, however, *spoh1* exudates from sucrose-grown seedlings induced significantly more prehaustoria in WT *P. japonicum*, compared to WT exudates, indicating a greater release of HIFs from *spoh1* roots (Fig 2C). *spoh1* exudates taken from plants grown in sucrose-containing media induced 70.7% of *Striga* seedlings to form prehaustoria (Fig 2D-F). These results demonstrate that the *spoh1* mutants grown in sucrose media secrete active HIFs.

**Fig. 2.**
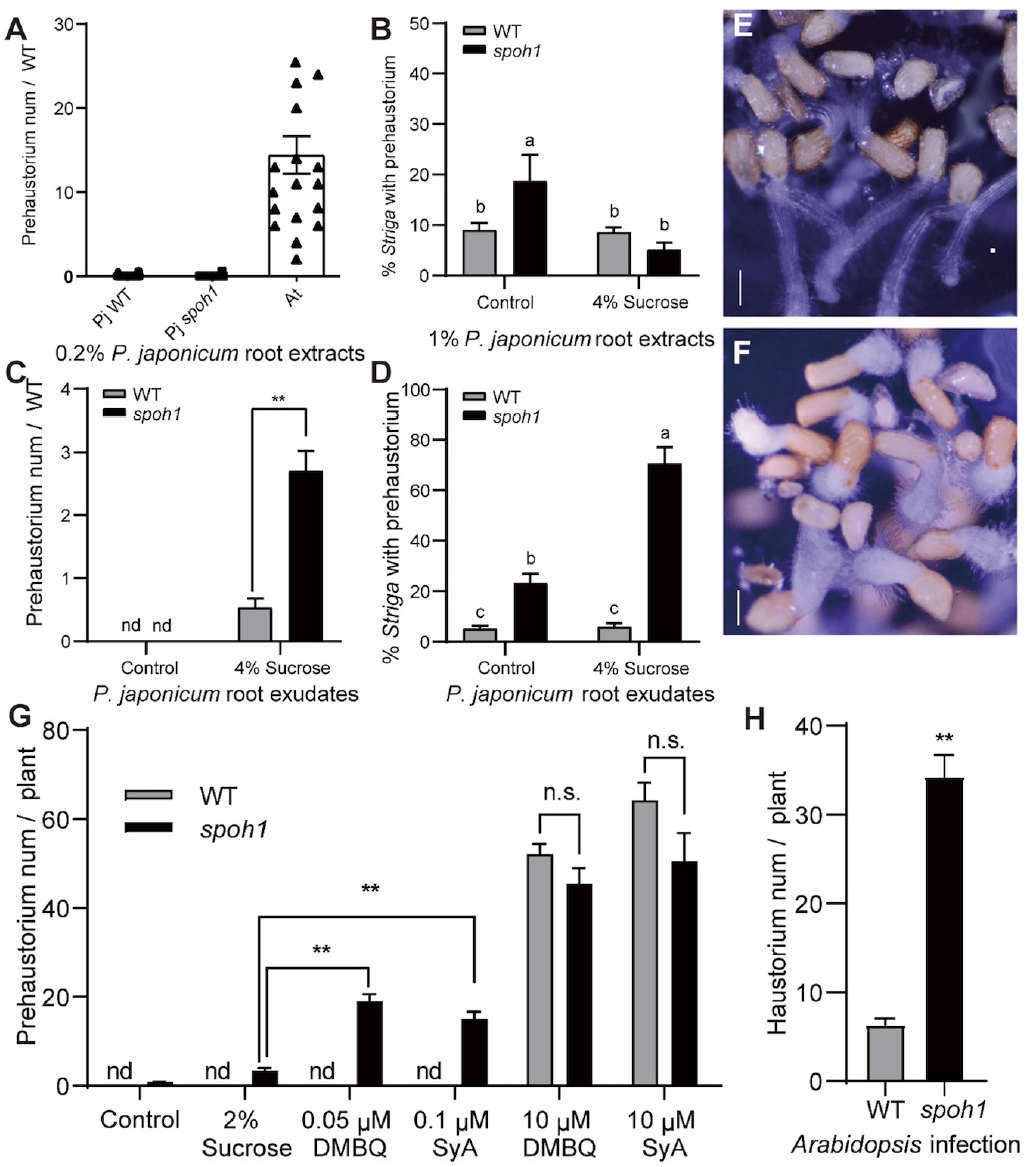
*spoh1* releases HIFs in root exudates and shows hypersensitivity to external HIFs. (**A**) *P. japonicum* prehaustorium number per plant induced by root extracts from WT and *spoh1* *P. japonicum* grown on 4% sucrose medium or wild-type *Arabidopsis*. Data represent mean ± SE from three representative experiments. n=30. (**B**) Prehaustorium formation rates of *S. hermonthica* induced by root extracts from WT and *spoh1 P. japonicum* grown on agar media with or without 4% sucrose-agar. Data represent mean ± SE from three representative experiments. Alphabet characters indicate significant difference detected by one-way ANOVA (*p*<0.05). (**C**) *P. japonicum* prehaustorium number per plant induced by root exudates of WT and *spoh1* grown with or without 4% sucrose. Experiments were repeated three times. n=30. Student’s t test, unpaired, two-tailed (\*\**p*< 0.01) (**D**) Prehaustorium formation rates of *S. hermonthica* induced by root exudates of WT and *spoh1* grown with or without 4% sucrose. n=3. One-way ANOVA. (**E, F**) *Striga* seedlings treated with root exudates of WT (left) and *spoh1*(right) grown with 4% sucrose. Scale bar: 200 μm. (**G**) Prehaustorium number of WT and *spoh1* in response to different concentration of HIFs (DMBQ, SyA), counted after 14-day treatments. Data are mean ± SE. from three representative experiments. n = 30. Student’s *t* test, unpaired, two-tailed (\*\**p*<0.01). (**H**) Haustorium number of WT and *spoh1* in Arabidopsis interaction. n = 15.

Although *spoh1* root exudates grown without sucrose did not show significant HIF activity (Fig 2C), *spoh1* nonetheless formed spontaneous prehaustoria under such conditions (Fig 1D-F, Fig 2G), indicating that *spoh1* exhibits higher sensitivity to HIFs than WT. To assess this, *spoh1* seedlings were treated with low concentrations of DMBQ (0.05 µM) and syringic acid (SyA) (0.1 µM), concentrations at which WT *P. japonicum* failed to form prehaustoria. Under these conditions, *spoh1* produced a significantly higher number of prehaustoria (Fig 2G), suggesting that *spoh1* is capable of responding to lower HIF concentration than WT. However, in the presence of 10 µM DMBQ or SyA, *spoh1* formed a comparable number of prehaustoria to WT, indicating that the maximum capacity for prehaustoria formation remains limited in *spoh1* (Fig 2G). Furthermore, *spoh1* formed significantly more prehaustoria with Arabidopsis (Fig 2H), indicating that *spoh1* is hypersensitive to both HIFs and host signals.

### The *spoh1* phenotype is caused by a mutation in a UDP-glucosyltransferase

To identify the *spoh1* gene, we sequenced F2 progeny genomes after backcrossing twice with WT, and examination of phenotypic segregation in F1 and F2 progenies. Most of the F1 progeny had the WT phenotype, with fewer than 10% displaying the *spoh1* phenotype (Table S1). The F2 progeny showed a 3:1 segregation ratio of WT to *spoh1* phenotypes, indicating that the *spoh1* phenotype is due to a single recessive mutation, although the SPOH1 function may be dose-dependent. The genome sequence of bulked *spoh1* phenotype F2 plants found two non-synonymous single nucleotide polymorphisms (SNPs) with frequencies exceeding 95%, emerging as the potential *spoh1* mutations (Data S1). The first candidate SNP was in the gene encoding a homolog of Arabidopsis GT72B1, a UDP-glucose-dependent glucosyltransferase, with a nonsense mutation at Lys 226 (Fig. 3A). The second candidate gene encodes a homolog of the Wiskott-Aldrich Syndrome (WAS) or WAS-like (WASL)-interacting family protein with an unknown function, bearing a Lys 109 to Glu missense mutation (Data S1).

**Fig. 3.**
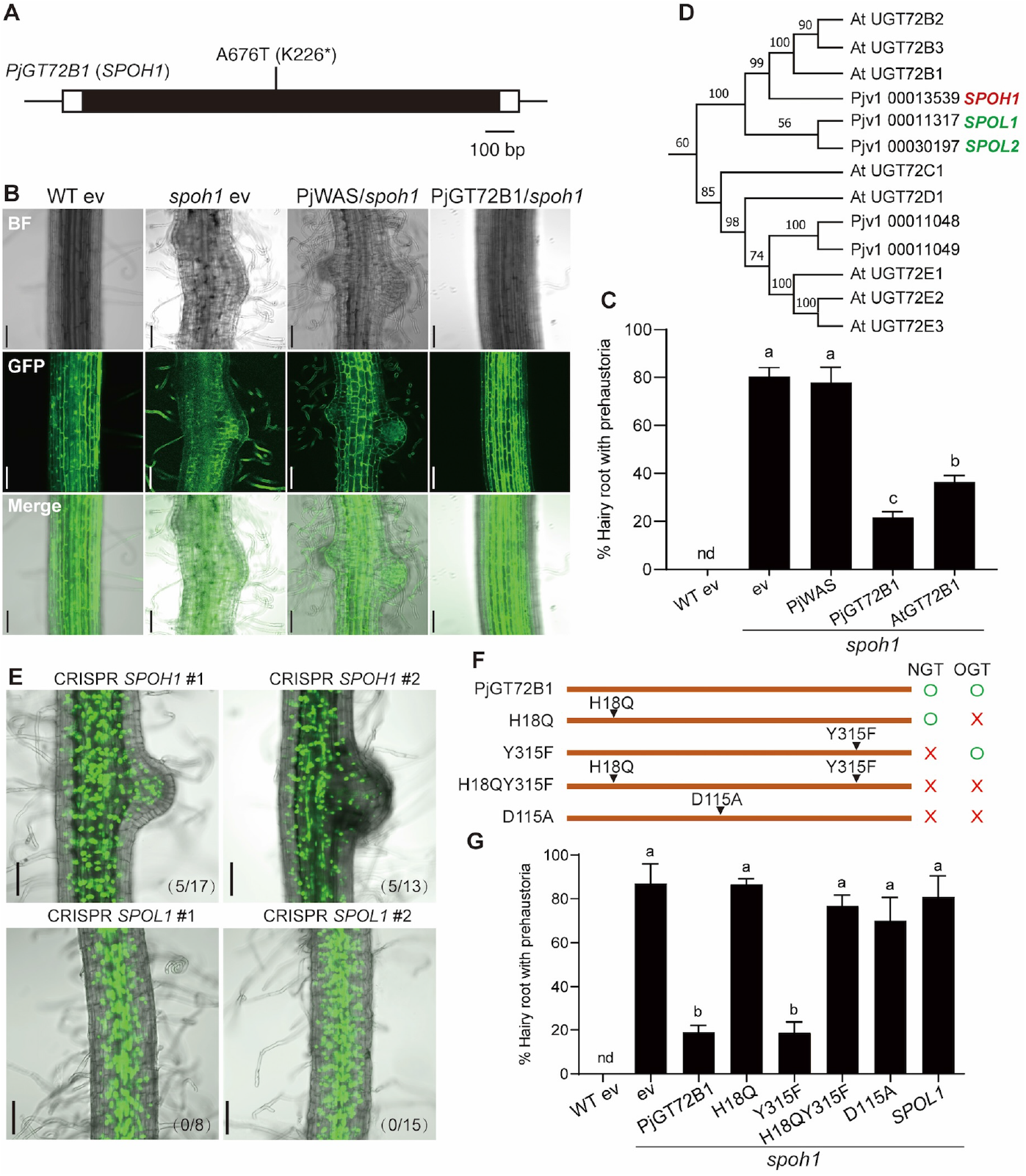
*spoh1* phenotype is caused by a mutation in GT72B1. (**A**) The gene structure of PjGT72B1. *spoh1* mutant possesses a non-sense mutation at the indicated position. Black box: coding sequence, White box: 5’ and 3’ UTRs. **(B)** Phenotypes of hairy roots transformed with empty vector (ev) or overexpressing *PjWAS* or *PjGT72B1*. The hairy roots with transformation marker GFP fluorescence were treated with 0.05 μM DMBQ for 1 week. Scale bar: 100 μm. **(C)** Percentage of hairy roots with prehaustoria after 7-day incubation on 0.05 μM DMBQ medium. Data represent mean ± SE from three representative experiments. n=27∼47. Significant difference was detected by one-way ANOVA (p < 0.05). (**D**) Phylogenetic tree of UGT72B subclade from *P. japonicum* and Arabidopsis UGTs. Numbers on the branch represent bootstrap value. (**E**) Phenotype of WT hairy roots transformed with CRISPR/Cas9 construct targeting to *SPOH1* and *SPOL1*. GFP-positive hairy roots were treated with 0.05 μM DMBQ for 2 weeks. Numbers at right bottom corner represent number of prehaustorium forming roots slash total observed hairy roots. Scale bar: 100 μm. (**F**) Schematic diagram of *PjGT72B1* mutants and their activities. (**G**) Percentage of hairy roots with prehaustoria induced by 0.05 μM DMBQ for 1 week. Data are mean ± SE from three representative experiments. n =16∼44. Significant difference was detected by one-way ANOVA(p<0.05).

Phenotypic complementation was evaluated in *spoh1* transgenic hairy roots overexpressing each candidate genes *PjGT72B1* (*pUB::PjGT72B1*) or *PjWAS* (*pUB::PjWAS*) with co-transformed GFP marker (Fig. 3B). WT hairy roots transformed with the empty vector produced very few prehaustoria, but nearly 80% of *spoh1* hairy roots with the empty vector formed prehaustoria, with an average of four per hairy root in response to 0.05 µM DMBQ for 7 days (Fig 3C). *spoh1* transgenic roots overexpressing *PjWAS* showed the similar percentage of prehaustorium-forming hairy roots and number of prehaustoria per hairy root as the empty vector control. In contrast, hairy roots overexpressing *PjGT72B1* had far fewer percentage and prehaustorium number than the control (Fig 3B, C, fig S4A). RT-qPCR confirmed the overexpression of *PjGT72B1* and *PjWAS* in the hairy roots, and importantly the expression levels of *PjGT72B1* correlated with the degree of phenotypic complementation (fig S4 B-C), indicating that the *spoh1* phenotype is attributable to a mutation in *PjGT72B1*.

The role of *PjGT72B1* was further confirmed by gene knock-out experiments. A BLAST search using the conserved PSPG box characteristic of UGT families identified 90 UGT-like sequences in the *P. japonicum* genome. A phylogenetic tree including both *P. japonicum* and Arabidopsis UGT sequences clustered into 14 distinct subfamilies (fig. S5)(*17*). The *P. japonicum* genome has UGTs in every subfamily (fig S5). Within the GT72B subclade of subfamily E, two additional genes were identified alongside Pjv1_00013539 (SPOH1): Pjv1_00011317 and Pjv1_00030197, which we have designated as SPOH1-like1 (*SPOL1*) and *SPOL2*, respectively (Fig3D, fig S8). Public RNA-seq data revealed that *SPOL2* is not expressed in roots. Therefore, we generated knock-out hairy roots for both *PjGT72B1* and *SPOL1* using CRISPR/Cas9 system in a WT background. Exogenous treatment with 0.05 µM DMBQ induced prehaustorium formation only in *PjGT72B1*-KO hairy roots, confirming *PjGT72B1* as the *SPOH1* gene (Fig 3E). Additionally, overexpression of *SPOL1* in the *spoh1* background failed to rescue the phenotype (Fig 3G, Fig S6 A-B), suggesting that *SPOH1* is uniquely responsible for suppressing spontaneous prehaustorium formation, and that *SPOL1* may have other functions.

To investigate functional conservation of GT72B1 across species, the Arabidopsis GT72B1 homolog (AtGT72B1) was overexpressed in *spoh1* hairy roots, resulting in significantly reduced prehaustorium formation compared to the *spoh1* empty vector control, albeit significantly higher than hairy roots overexpressing *PjGT72B1* (Fig 3C). RT-qPCR analysis confirmed the overexpression of *AtGT72B1* in hairy roots, indicating that the partial complementation was not due to insufficient gene expression (fig S4D). These results suggest that while *AtGT72B1* shares a similar function with *PjGT72B1*, its activity is not fully equivalent.

### *O*-Glucosyltransferase activity is essential for *spoh1* complementation

AtGT72B1 is a bifunctional enzyme with both *O*-glucosyltransferase (OGT) activity, catalyzing the glycosylation of the hydroxyl groups in phenols, and *N*-glycosyltransferase (NGT) activity, which catalyze the glycosylation of the amine groups in anilines (*18*). To investigate whether PjGT72B1 is also bifunctional, we generated glutathione *S*-transferase (GST)-fused PjGT72B1 and AtGT72B1 in *E. coli*, and conducted *in vitro* glycosylation assays (fig. S7). We used 2,4,5-trichlorophenol (TCP) and 3,4-dichloroaniline (DCA) as standard substrates for the OGT and NGT activities, respectively (fig. S8A,B). Additional HPLC product peaks were evident in PjGT72B1 reactions with both TCP and DCA (fig. S8C and D), which corresponded to the glucose-conjugated forms of the substrates, as confirmed by LC-MS (Fig. S8E,F).

To assess whether both OGT and NGT activities are required for *SPOH1*’s function in suppressing spontaneous prehaustorium formation, we generated modified PjGT72B1 proteins that retain either OGT or NGT activity, or which had lost both activities. Sequence alignments with AtGT72B1 indicated important residues for each enzyme activity (fig. S9) (*19*), and we generated the mutants with NGT activity only (H18Q), OGT activity only (Y315F), loss of both activities (H18QY315F), and loss of catalytic activity (D115A) (Fig. 3F). The predicted enzyme activities of each modified PjGT72B1 protein were confirmed by *in vitro* glycosylation assays (fig. S7, S8). These mutants were then expressed in *spoh1* to assess phenotypic complementation. Neither the H18QY315F double mutant nor D115A rescued the *spoh1* phenotype, underscoring the necessity of enzyme activity for *SPOH1* function (Fig. 3G, fig. S6). In contrast, the Y315F mutant successfully complemented the *spoh1* phenotype, but the H18Q mutant did not, indicating that OGT activity, rather than NGT activity, is responsible for suppressing spontaneous prehaustorium formation (Fig. 3G, fig. S6).

### PjGT72B1 modulates lignin biosynthesis and suppresses prehaustorium induction

To understand the *in planta* function of PjGT72B1, the *spoh1* mutants grown with or without 2% sucrose were subjected to RNA-seq analysis (Fig. 4A, fig.S10A,B, Data S2-5). Among the 133 genes consistently and significantly upregulated in *spoh1* compared to WT were several known to be involved in prehaustorium formation, including *PjQR2* (*20*), a homolog of *TvPirin* (*21*), and the early HIF responsive genes (*22, 23*), confirming that a set of prehaustorium genes are activated in *spoh1*(Fig. 4B and C).

**Fig. 4.**
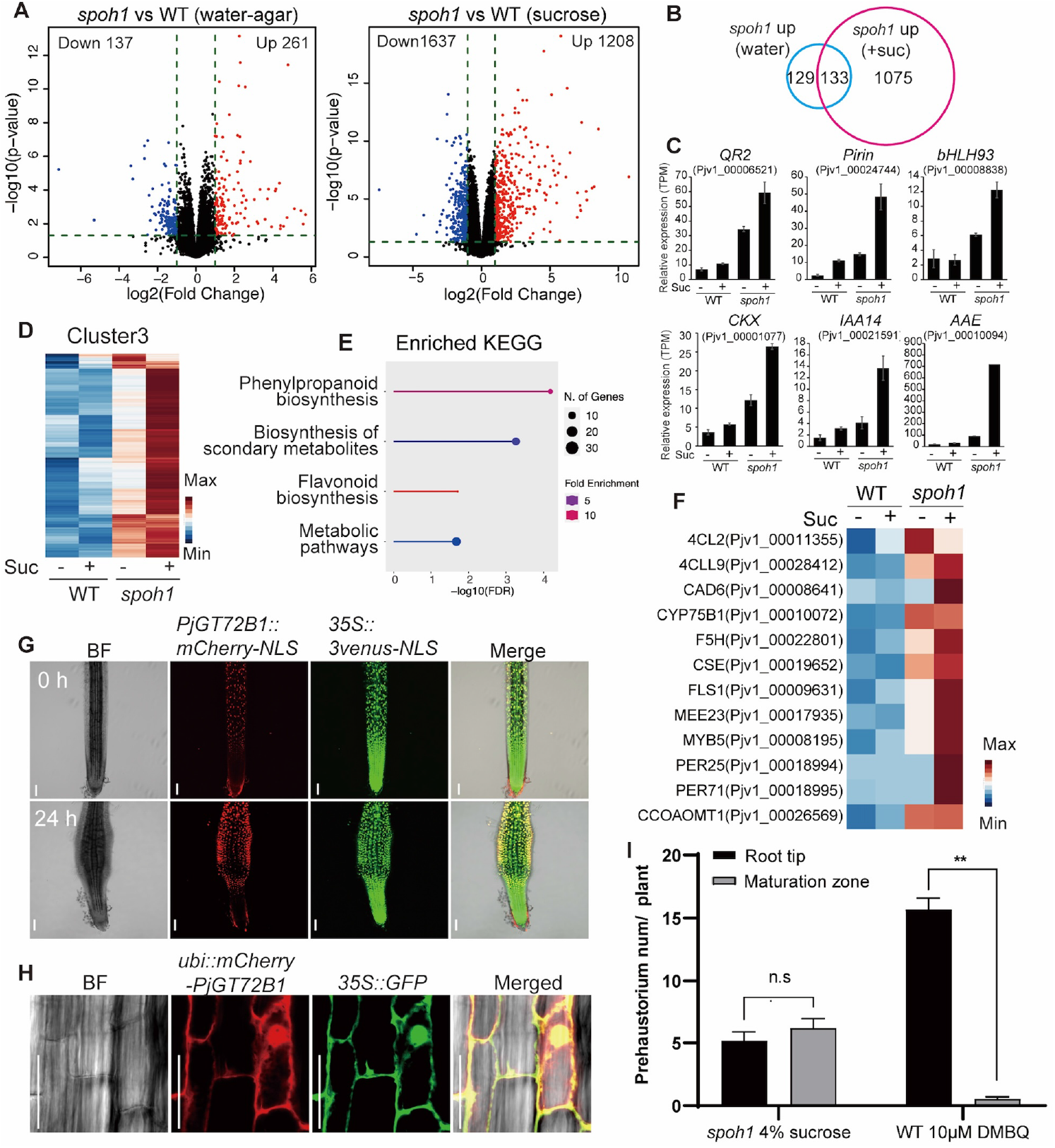
*PjGT72B1* modulates endogenous lignin biosynthesis in the maturation zone and suppresses self-prehaustorium formation. **(A)** Volcano plots of RNA-sequencing results comparing WT and *spoh1* with (water-agar) or without 2% sucrose (sucrose) condition. Up and down-regulated genes in *spoh1*were highlighted with red and blue colors, respectively. **(B)** Venn diagram from (A). **(C)** Expression patterns of known marker genes for early HIF response, detected as commonly upregulated genes in *spoh1*. **(D)** A heatmap of Cluster3 genes from SOM clustering (fig. S13E). **(E)** Enrichment analysis of KEGG pathway in Cluster3 genes. **(F)** Expression patterns of lignin biosynthesis related genes involved in Cluster3. **(G)** PjGT72B1 expression pattern before (0h) or after (24h) prehaustorium induction by 0.4% Arabidopsis root extracts. PjGT72B1 promoter activity is monitored by mCherry fluorescence, and Venus-NLS was used as a transformation marker. Scale bar: 100 μm. **(H)** Subcellular localization of PjGT72B1 in *P. japonicum* hairy roots was detected as protein fusion with mCherry. 35S-GFP serves as a control for cytosol and nuclei localization. Scale bar: 50 μm. **(I)** Prehaustorium formation sites of WT treated with 10 µM DMBQ and *spoh1* incubated on 4% sucrose media. Newly formed prehaustorium at root tip or maturation zone were counted every 24h for 13 days. Data are mean ± SE from three representative experiments; n = 30. Student’s t test, unpaired, two-tailed (\*\**p* < 0.01).

To further dissect the molecular effects of *spoh1*, we classified 6,968 differentially expressed genes across all sample combinations into 12 clusters using self-organized mapping (SOM) (fig. S10C-E, Data S6). The 263 genes in Cluster 3 had higher expression in *spoh1* than in WT, and their expression was further increased by the addition of sucrose addition (Fig 4D). Gene ontology enrichment and network analyses indicated significant enrichment of KEGG pathways related to the phenylpropanoid metabolic and biosynthesis processes, which include the lignin biosynthesis pathway (Fig 4E, fig. S11, fig. S12). Importantly, Cluster 3 encompasses genes encoding lignin biosynthesis enzymes, such as 4CL, CAD, F5H and peroxidases, along with a homolog of the MYB5 transcription factor, suggesting that the *spoh1* mutation activates lignin biosynthesis, a process further stimulated by sucrose (Fig 4F). This is in line with the previous observation in the Arabidopsis *atgt72b1* mutant, which overaccumulates lignin and upregulates lignin biosynthesis genes (*24*). Because endogenous lignin biosynthesis facilitates the production of HIFs in hosts and prehaustorium formation in *S. hermonthica* (*10*), the upregulation of lignin biosynthesis in *spoh1* may contribute to g endogenous HIF production and hypersensitivity to external HIFs.

The promoter activity of *PjGT72B1*, visualized by *PjGT72B1p-mCherry-NLS* construct, was detected in the maturation zone of roots, whereas no signal was observed in the meristematic zone regardless of HIF application (Fig 4G, fig. S13A, B). In the elongation zone, where prehaustoria generally form in response to HIFs, a weaker mCherry signal was detected, which intensified as prehaustorium developed (Fig 4G, fig. S13A, B). During host invasion, a strong mCherry signal was observed at the interface with host tissues, particularly in the intrusive cells at the haustorium apex (fig. S14), suggesting a potential role for *PjGT72B1* during host invasion. Localization studies of a constitutively expressed fusion protein *pUB::mCherry-PjGT72B1*, which complemented the *spoh1* phenotype (Fig 4H, fig S15), indicated that PjGT72B1is localized to both the cytosol and nuclei. This suggests that PjGT72B1 glycosylates intracellular phenolic compounds mainly in the root maturation zone.

Because lignin synthesis is more active in the root maturation zone than in the meristematic zone, it is likely that the primary location for PjGT72B1 glucosylation of lignin monomers or their precursors for inactivation of endogenous HIFS is within the maturation zone. Supporting this, we observed that *spoh1* tends to form more prehaustoria in the maturation zone. About half of *spoh1* spontaneous prehaustoria formed in the maturation zone, compared to 98% of HIF-induced prehaustoria in WT forming in the elongation zone (Fig 4I, fig. S16). This suggests that the PjGT72B1 mutation allows *P. japonicum* to respond to its own lignin-related compounds, thus triggering spontaneous prehaustorium formation in the maturation zone.

### PjGT72B1 catalyzes HIF glucosylation with different substrate specificity from AtGT72B1

Given that *spoh1* produces more HIFs in its root exudates than WT, and that phenolic glucosylation is crucial for suppressing spontaneous prehaustorium formation, we hypothesized that PjGT72B1 mediates the glucosylation of HIFs produced by *P. japonicum*. The glucosylation would thereby prevent self-induced prehaustorium formation by inactivating endogenous HIFs. To test this model, we assayed *in vitro* enzyme activity on vanillin, SyA, FerA and sinapic acid (SinA), as potential substrates (Fig 5A,B, fig. S17A,B). WT PjGT72B1, but not catalytic inactive D115A, glucosylated all of these substrates, shown by new LC-MS peaks corresponding to glucosylated substrates (Fig 5, fig S17). Interestingly, AtGT72B1 glucosylated the G-type HIFs, vanillin and FerA, but failed to act on the S-type HIFs, SyA and SinA (Fig 5A,B, fig. S17A,B). This is notable because the S-type HIFs generally exhibits higher activity than the G-type HIFs (*10*). This difference in substrate specificity could explain why AtGT72B1 only partially complements the *spoh1* phenotype; the S-type HIFs likely remain active for prehaustorium induction.

**Fig. 5.**
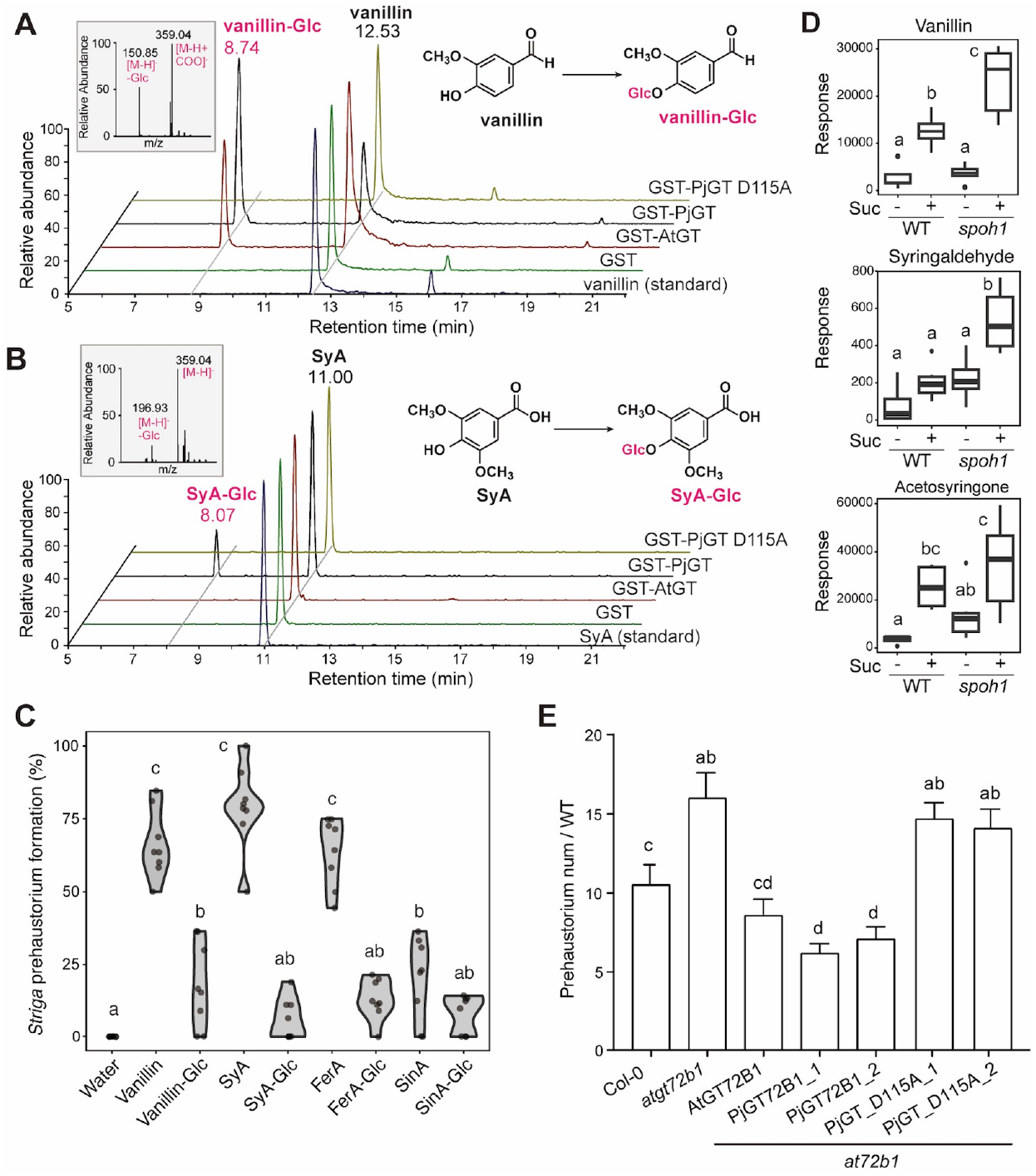
PjGT72B1 regulates HIF activity in the parasite and the host, by glucosylation and subsequent inactivation of HIFs. **(A, B)** *In vitro* glucose transfer to vanillin (A) and SyA (B) by GT72B1 analyzed by LC-MS. GST only and GST-PjGT72B1 D115A mutant (PjGT D115A) were used as negative control. STD indicates chromatogram of the standard substrate. Blue and red arrows indicate substrate and product peaks, respectively. Small insets show MS peaks of the products. (**C**) Prehaustorium formation rates in *S. hermonthica* treated with various HIFs (vanillin, SyA, FerA, and SinA) and their corresponding glucosides (vanillin-Glc, SyA-Glc, FerA-Glc, SinA-Glc). n = 8. Different alphabets indicate significant difference detected by Tukey HDS test (*p* < 0.05). **(D)** Relative amounts of HIFs in root exudates of WT and *spoh1* grown with or without sucrose were measured by LC-MS. **(E)** Prehaustorium number per plant in WT *P. japonicum* treated by root exudates of Arabidopsis Col-0 and *atgt72b1* mutant overexpressing *AtGT72B1* and *PjGT72B1* as well as the *PjGT72B1* D115A mutant. n=30. Alphabet characters represent significantly different value detected by one-way ANOVA (p<0.05). with 0.05 μM DMBQ for 2 weeks. Numbers at right bottom corner represent number of prehaustorium forming roots slash total observed hairy roots. Scale bar: 100 μm. (**F**) Schematic diagram of *PjGT72B1* mutants and their activities. (**G**) Percentage of hairy roots with prehaustoria induced by 0.05 μM DMBQ for 1 week. Data are mean ± SE from three representative experiments. n =16∼44. Significant difference was detected by one-way ANOVA(p<0.05).

To further validate the inactivation of HIFs by glucosylation, the prehaustorium induction activity of the HIF glucosides, vanillin-Glc, FerA-Glc and SyA-Glc, were tested on *S. hermonthica*. While the unmodified forms of vanillin, SyA and FerA demonstrated robust HIF activity, their glucosylated forms, vanillin-Glc, SyA-Glc and FerA-Glc, showed significantly reduced HIF activities (Fig 5C). Consistent with a previous study(*10*), SinA exhibited weak HIF activity, and its glucoside (SinA-Glc) had correspondingly low HIF activity (Fig. 5C). Overall, these results indicate that HIF glucosylation nearly eliminates prehaustorium-induction activity, strongly supporting the hypothesis that PjGT72B1 modulates prehaustorium induction through HIF glucosylation.

### GT72B1 regulates the HIF production of host plants

To confirm that *spoh1* exudes more HIFs compared to WT *P. japonicum*, HIFs in the root exudates were quantified. When grown in 4% sucrose, *spoh1* indeed exuded larger amounts of vanillin, syringaldehyde and acetosyringone, than WT (Fig. 5D). Other HIFs, including DMBQ, SyA, FerA and vanillic acid were not higher in *spoh1* root exudates than in WT (fig. S18). The specific contribution of each of these HIFs to prehaustorium induction remains to be elucidated.

GT72B1 may also contribute to the glucosylation of host-derived HIFs, thereby modulating prehaustorium induction activity. To test this possibility, we used the root exudates of the Arabidopsis *atgt72b1* T-DNA mutant, which induced significantly more prehaustoria in WT *P. japonicum* compared to Col-0, indicating that *AtGT72B1* normally suppresses endogenous HIF activity in the host plant (Fig 5E). Overexpression of *AtGT72B1* rescued the *atgt72b1* phenotype, resulting in induction of the comparable number of prehaustoria with Col-0, while overexpression of *PjGT72B1* displayed lower prehaustorium induction activity compared to Col-0 (Fig 5E). These results suggest that the broader spectrum of HIF substrates of *PjGT72B1* confers stronger HIF suppression than *AtGT72B1*.

## Conclusions

Here we have demonstrated that the glucosylation of phenolic HIFs by GT72B1 plays a crucial role in kin avoidance in parasitic plants. Specifically, PjGT72B1 suppresses formation of self-prehaustoria in *P. japonicum* by inactivating HIFs. While PjGT72B1 can glucosylate both G-type and S-type HIFs, its host counterpart AtGT72B1 selectively glucosylates only G-types, thereby allowing the exudation of active HIFs. This difference in substrate specificity may underpin kin avoidance mechanisms in parasitic plant species (fig. S19). Glucosylated lignin precursors serve as mobile and storage forms, thereby facilitating the precise modulation of lignin biosynthesis (*25*). Our findings demonstrate a novel role for lignin monomer glucosylation, regulating their signaling activity for plant kin avoidance. The *spoh1* mutation results in increased lignin biosynthesis gene expression and endogenous HIF production, enhancing spontaneous prehaustorium formation. However, in the absence of sucrose, the prehaustorium induction by *spoh1* root exudate remains lower than that of the host plant, implying that additional mechanisms restrain HIF exudation in parasitic plants. Moreover, the growth defects observed in both Arabidopsis *atgt72b1* (*24*) and *spoh1* suggest the involvement of these glucosyltransferases in plant growth and development. Together, our findings underscore the kin avoidance mechanism in parasitic plants, and the regulation of HIF production in their host. This knowledge may ultimately be applied to develop crop plants with lower HIF production to improve the control of these devastating parasitic weeds.

## Acknowledgments

The authors thank Prof. Abdelbagi M Ali and late Prof. A. G. Babiker for providing *S. hermonthica* seeds, and late Emer. Prof. Kenji Mori for providing strigol. We thank Mrs. Maki Nozaki and Yuko Yoshimura for their technical assistance. LX and MC were supported by the MEXT international student fellowship program. A part of this study was conducted using the facilities at the DASH/FBAS, RISH, Kyoto University, and the NMR spectrometer at JURC, ICR, Kyoto University. This work was partly supported by the NAIST Life Science Collaboration Center (LiSCo). The identification of the mutant gene was partly supported by JSPS KAKENHI Grant Number JP22H04925 (PAGS).

## Funding

Japan Society for the Promotion of Science KAKENHI grant JP20H05909 (SY, KS)

Japan Society for the Promotion of Science KAKENHI grant JP21H02506 (SY, KS)

Japan Society for the Promotion of Science KAKENHI grant JP24K01827 (YT)

Japan Society for the Promotion of Science KAKENHI grant JP22H02648(TT)

National Natural Science Foundation of China grant number 32370317 (SC)

The Chinese Academy of Sciences “Introducing talents funds of Kunming Institute of Botany” (SC)

Yunnan Revitalization Talent Support Program “Young Talent” Project (SC)

JST PRESTO grant number JPMJPR194D (SY) Mitsubishi Foundation (SY)

## Author contributions

Conceptualization: SC, SBS, KS, SY

Methodology: LX, SC, YT, MS, HB, MW, TT, SI

Investigation: LX, SC, SBS, MT, SI, XB, SS, MW, MC, MS, HB, TT,SY

Visualization: LX, MC, SY

Funding acquisition: SC, YT, TT, KS, SY

Project administration: SC, KS, SY

Supervision: SC, YT, HB, KS, SY

Writing – original draft: LX, SY

Writing – review & editing: LX, SC, SI, YT, HB, KS, SY

## Competing interests

Authors declare that they have no competing interests.

## Data and materials availability

The genome sequence data and RNA sequencing data used in this manuscript were deposited to DDBJ with accession number PRJNA1202847.

## List of Supplementary Materials

Materials and Methods

Supplementary Text

Figs. S1 to S19

Tables S1 to S2

Data S1 to S6

## Notes

### Competing Interest Statement

The authors have declared no competing interest.

## References

1. S. Runo, E. K. Kuria, Habits of a highly successful cereal killer, Striga. PLoS Pathog. 14, e1006731 (2018).

2. E. Pennisi, Armed and dangerous. Science. 327, 804–5 (2010).

3. S. Yoshida, S. Cui, Y. Ichihashi, K. Shirasu, The haustorium, a specialized invasive organ in parasitic plants. Annu. Rev. Plant Biol. 67, 643–667 (2016).

4. J. I. Yoder, A species-specific recognition system directs haustorium development in the parasitic plant Triphysaria (Scrophulariaceae). Planta. 202, 407–413 (1997).

5. J. M. Mutuku, S. Cui, S. Yoshida, K. Shirasu, Orobanchaceae parasite–host interactions. New Phytol. 230, 46–59 (2020).

6. K. M. Furuta, L. Xiang, S. Cui, S. Yoshida, Molecular dissection of haustorium development in Orobanchaceae parasitic plants. Plant Physiol. 186, 1424–1434 (2021).

7. H. Albrecht, J. I. Yoder, D. A. Phillips, Flavonoids promote haustoria formation in the root parasite triphysaria versicolor. Plant Physiol. 119, 585–592 (1999).

8. V. Goyet, S. Wada, S. Cui, T. Wakatake, K. Shirasu, G. Montiel, P. Simier, S. Yoshida, Haustorium inducing factors for parasitic Orobanchaceae. Front. Plant Sci. 10, 1056 (2019).

9. C. R. Clarke, M. P. Timko, J. I. Yoder, M. J. Axtell, J. H. Westwood, Molecular Dialog Between Parasitic Plants and Their Hosts. Annu. Rev. Phytopathol. 57, 1–21 (2019).

10. S. Cui, S. Wada, Y. Tobimatsu, Y. Takeda, S. B. Saucet, T. Takano, T. Umezawa, K. Shirasu, S. Yoshida, Host lignin composition affects haustorium induction in the parasitic plants Phtheirospermum japonicum and Striga hermonthica. New Phytol. 218, 710–723 (2018).

11. S. B. Saucet, K. Shirasu, Molecular Parasitic Plant–Host Interactions. PLoS Pathog. 12, 1–6 (2016).

12. S. Yoshida, K. Shirasu, Multiple layers of incompatibility to the parasitic witchweed, Striga hermonthica. New Phytol. 183, 180–189 (2009).

13. Y. Wang, M. Murdock, S. W. T. Lai, D. B. Steele, J. I. Yoder, Kin recognition in the parasitic plant Triphysaria versicolor is mediated through root exudates. Front. Plant Sci. 11, 1–11 (2020).

14. S. Cui, T. Wakatake, K. Hashimoto, S. Saucet, K. Toyooka, S. Yoshida, K. Shirasu, Haustorial hairs are specialized root hairs that support parasitism in the facultative parasitic plant, Phtheirospermum japonicum. Plant Physiol. 170, 1492–1503 (2016).

15. L. Xiang, Y. Li, X. Sui, A. Li, Fast and abundant in vitro spontaneous haustorium formation in root hemiparasitic plant Pedicularis kansuensis Maxim. (Orobanchaceae). Plant Divers. 40, 226–231 (2018).

16. J. Yoon, L. H. Cho, W. Tun, J. S. Jeon, G. An, Sucrose signaling in higher plants. Plant Sci. 302, 110703 (2021).

17. Y. Li, S. Baldauf, E. K. Lim, D. J. Bowles, Phylogenetic Analysis of the UDP-glycosyltransferase Multigene Family of Arabidopsis thaliana. J. Biol. Chem. 276, 4338– 4343 (2001).

18. M. Brazier-Hicks, R. Edwards, Functional importance of the family 1 glucosyltransferase UGT72B1 in the metabolism of xenobiotics in Arabidopsis thaliana. Plant J. 42, 556–566 (2005).

19. M. Brazier-Hicks, W. A. Offen, M. C. Gershater, T. J. Revett, E. K. Lim, D. J. Bowles, G. J. Davies, R. Edwards, Characterization and engineering of the bifunctional N- and O-glucosyltransferase involved in xenobiotic metabolism in plants. Proc. Natl. Acad. Sci. U. S. A. 104, 20238–20243 (2007).

20. J. K. Ishida, S. Yoshida, K. Shirasu, Quinone oxidoreductase 2 is involved in haustorium development of the parasitic plant Phtheirospermum japonicum. Plant Signal. Behav. 12, 1–4 (2017).

21. P. C. G. Bandaranayake, A. Tomilov, N. B. Tomilova, Q. a Ngo, N. Wickett, C. W. dePamphilis, J. I. Yoder, The TvPirin gene is necessary for haustorium development in the parasitic plant Triphysaria versicolor. Plant Physiol. 158, 1046–53 (2012).

22. J. K. Ishida, T. Wakatake, S. Yoshida, Y. Takebayashi, H. Kasahara, E. Wafula, C. W. DePamphilis, S. Namba, K. Shirasu, Local auxin biosynthesis mediated by a YUCCA flavin monooxygenase regulates haustorium development in the parasitic plant Phtheirospermum japonicum. Plant Cell. 28, 1795–1814 (2016).

23. N. Aoki, S. Cui, C. Ito, K. Kumaishi, S. Kobori, Y. Ichihashi, S. Yoshida, Phenolic signals for prehaustorium formation in Striga hermonthica. Front. Plant Sci. 13, 1–15 (2022).

24. J. S. Lin, X. X. Huang, Q. Li, Y. Cao, Y. Bao, X. F. Meng, Y. J. Li, C. Fu, B. K. Hou, UDP-glycosyltransferase 72B1 catalyzes the glucose conjugation of monolignols and is essential for the normal cell wall lignification in Arabidopsis thaliana. Plant J. 88, 26–42 (2016).

25. T. Tsuyama, R. Kawai, N. Shitan, T. Matoh, J. Sugiyama, A. Yoshinaga, K. Takabe, M. Fujita, K. Yazaki, Proton-dependent coniferin transport, a common major transport event in differentiating xylem tissue of woody plants. Plant Physiol. 162, 918–926 (2013).

26. S. Cui, T. Kubota, T. Nishiyama, K. Juliane, S. Shigenobu, T. F. Shibata, A. Toyoda, M. Hasebe, K. Shirasu, S. Yoshida, Ethylene signaling mediates host invasion by parasitic plants. Sci. Adv. 6, eabc2385 (2020).

27. S.Y. Songkui Cui, Abdelbagi M.A. Ghanim, in Mutation Breeding and Efficiency Enhancing Technologies for Resistance to Striga in Cereals, A. M. A. G. S. Sivasankar, Ed. (Springer, 2024), pp. 145–160.

28. T. Maekawa, M. Kusakabe, Y. Shimoda, S. Sato, S. Tabata, Y. Murooka, M. Hayashi, Polyubiquitin promoter-based binary vectors for overexpression and gene silencing in Lotus japonicus. Mol. Plant. Microbe. Interact. 21, 375–82 (2008).

29. C. Engler, M. Youles, R. Gruetzner, T. M. Ehnert, S. Werner, J. D. G. Jones, N. J. Patron, S. Marillonnet, A Golden Gate modular cloning toolbox for plants. ACS Synth. Biol. 3, 839–843 (2014).

30. J. K. Ishida, S. Yoshida, M. Ito, S. Namba, K. Shirasu, Agrobacterium rhizogenes-mediated transformation of the parasitic plant Phtheirospermum japonicum. PLoS One. 6, e25802 (2011).

31. A. Greifenhagen, I. Braunstein, J. Pfannstiel, S. Yoshida, K. Shirasu, A. Schaller, T. Spallek, The Phtheirospermum japonicum isopentenyltransferase PjIPT1a regulates host cytokinin responses in Arabidopsis. New Phytol. 232, 1582–1590 (2021).

32. A. M. Bolger, M. Lohse, B. Usadel, Trimmomatic: A flexible trimmer for Illumina sequence data. Bioinformatics. 30, 2114–2120 (2014).

33. D. Kim, J. M. Paggi, C. Park, C. Bennett, S. L. Salzberg, Graph-based genome alignment and genotyping with HISAT2 and HISAT-genotype. Nat. Biotechnol. 37, 907–915 (2019).

34. S. Yoshida, S. Kim, E. K. Wafula, J. Tanskanen, Y. M. Kim, L. Honaas, Z. Yang, T. Spallek, C. E. Conn, Y. Ichihashi, K. Cheong, S. Cui, J. P. Der, H. Gundlach, Y. Jiao, C. Hori, J. K. Ishida, H. Kasahara, T. Kiba, M. S. Kim, N. Koo, A. Laohavisit, Y. H. Lee, S. Lumba, P. McCourt, J. C. Mortimer, J. M. Mutuku, T. Nomura, Y. Sasaki-Sekimoto, Y. Seto, Y. Wang, T. Wakatake, H. Sakakibara, T. Demura, S. Yamaguchi, K. Yoneyama, R. ichiroh Manabe, D. C. Nelson, A. H. Schulman, M. P. Timko, C. W. DePamphilis, D. Choi, K. Shirasu, Genome sequence of Striga asiatica provides insight into the evolution of plant parasitism. Curr. Biol. 29, 3041–3052 (2019).

35. R. Wehrens, L. M. C. Buydens, Self-and super-organizing maps in R: the Kohonen package. J. Stat. Softw. 21, 19 (2007).

36. S. X. Ge, D. Jung, D. Jung, R. Yao, ShinyGO: A graphical gene-set enrichment tool for animals and plants. Bioinformatics. 36, 2628–2629 (2020).

37. D. Szklarczyk, A. Franceschini, S. Wyder, K. Forslund, D. Heller, J. Huerta-Cepas, M. Simonovic, A. Roth, A. Santos, K. P. Tsafou, M. Kuhn, P. Bork, L. J. Jensen, C. Von Mering, STRING v10: Protein-protein interaction networks, integrated over the tree of life. Nucleic Acids Res. 43, D447–D452 (2015).

38. Y. Miyagawa, T. Mizukami, H. Kamitakahara, T. Takano, Synthesis and fundamental HSQC NMR data of monolignol β-glycosides, dihydromonolignol β-glycosides and p-hydroxybenzaldehyde derivative β-glycosides for the analysis of phenyl glycoside type lignin-carbohydrate complexes (LCCs). 68, 747–760 (2014).

39. D. Kawa, B. Thiombiano, M. Z. Shimels, T. Taylor, A. Walmsley, H. E. Vahldick, D. Rybka, M. F. A. Leite, Z. Musa, A. Bucksch, F. Dini-Andreote, M. Schilder, A. J. Chen, J. Daksa, D.W. Etalo, T. Tessema, E. E. Kuramae, J. M. Raaijmakers, H. Bouwmeester, S.M. Brady, The soil microbiome modulates the sorghum root metabolome and cellular traits with a concomitant reduction of Striga infection. Cell Rep. 43, 113971 (2024).

40. K. Floková, M. Shimels, B. Andreo Jimenez, N. Bardaro, M. Strnad, O. Novák, H. J. Bouwmeester, An improved strategy to analyse strigolactones in complex sample matrices using UHPLC-MS/MS. Plant Methods. 16, 1–17 (2020).

