## Supplemental Materials for "Glucosylation of Endogenous Haustorium-inducing factors Underpins Kin Avoidance in Parasitic Plants"

**The PDF file includes:**

Materials and Methods
Figs. S1 to S19
Tables S1 to S2

**Other Supplementary Materials for this manuscript include the following:**

Data S1 to S6

### Materials and Methods

Plant materials and growth conditions

*Phtheirospermum japonicum* (Thunb.) Kanitz ecotype Okayama was used as the wild type (WT). *Arabidopsis thaliana* Columbia ecotype (Col-0) served as host plants. The Arabidopsis AtGT72B1 homozygous mutant (cs25067) was obtained from the Arabidopsis Resource Center (ABRC). *Striga hermonthica* (Delile) Benth. seeds were kindly provided by the late Professor Abdel Babikar and Professor Abdelbagi M Ali Gahmin.

*P. japonicum* seeds were surface sterilized with a 10% commercial bleach (approx. 6% sodium hypochlorite; Kao, Tokyo, Japan) for 8 min, washed five times with sterile water, and then sowed on half-strength Murashige and Skoog medium (1/2 MS) with 1% sucrose and 0.8% agar. Arabidopsis seeds were surface sterilized using a similar procedure, except with 5% commercial bleach. Both *P. japonicum* and Arabidopsis seeds were stratified for 1-2 days at 4°C in the dark before being transferred to a growth chamber. *P. japonicum* seeds were germinated at 25 ºC in the dark for 3 d, and then grown at 25 ºC under long-day conditions (16 h light / 8 h dark). Arabidopsis seeds were grown at 22 ºC under long-day conditions. For soil experiments, *P. japonicum* plants were grown at 23 ºC under long-day conditions for vegetative growth, followed by 25 ºC under short-day conditions (8 h light / 16 h dark) for reproductive growth.

*S. hermonthica* seeds were surface sterilized with a 20% bleach for 8 min, washed with sterile water for at least 5 times, and placed on glass fiber filter paper (Whatman GF/A) moistened with 10 ml sterile water. Sterilized seeds were incubated in the dark at 25 ºC for at least 5 days. Germination was induced by applying 10 nM strigol.

*spoh1* mutant isolation and backcrossing

Seeds from 0.3% EMS-mutagenized M2 progeny (*14*) were cultured in water-containing 96-well plates under 25 ºC long-day conditions with gentle shaking for 1 week. Under these conditions, WT seedlings produce long, thin root. WT seedlings incubated with 10 μM DMBQ produced short, curved roots with bump-like prehaustoria structures. Mutants displaying short, curved roots and prehaustoria after seven days of incubation in water were selected. Progeny seeds from the first screening were screened again, where 15 seedlings were incubated in water in 24-well plates for 3 days. The presumptive *spoh1* mutant was then backcrossed twice with WT to generate BC2F1 and BC2F2 progeny. *spoh1* BC2F3 progenies exhibiting the *spoh1* phenotype on sucrose medium were selected for further experiments.

<u>Quantification of spontaneous and HIF-induced prehaustoria</u>

Seven-day old WT and *spoh1* seedlings were transferred to 0.8% water agar medium for 3 days, after which they were transferred to 0.8% agar medium containing sucrose, glucose, fructose, galactose, maltose, raffinose, sorbitol, mannitol, or HIFs and oriented vertically. For sucrose treatment, sucrose was added to the 0.8% agar medium before autoclaving. The other sugars and sugar alcohols were dissolved in sterile water to make 40% (w/v) stocks, filter-sterilized, and then added to autoclaved 1% agar medium. Spontaneous prehaustoria were counted after 14 days incubation, unless otherwise indicated. For HIF treatments, DMBQ (Sigma-Aldrich) or syringic acid (SyA, Sigma-Aldrich), dissolved in DMSO, were added to autoclaved 0.8% agar medium at the indicated final concentration. Prehaustoria were counted after 14 days of induction, unless otherwise indicated.

<u>Observation of spontaneous prehaustoria formation site in *spoh1*</u>

Seven-day-old *spoh1* seedlings were transferred to 0.8% water agar for starvation for 3 days and then to 4% sucrose medium. The sites of newly-formed spontaneous prehaustoria were marked and recorded every 24 h over a 14-day observation period. Seven-day old WT seedlings were treated similarly, except that 10 μM DMBQ was used instead of 4% sucrose and the newly-formed prehaustoria were counted every 24 h for 14 days.

<u>Arabidopsis infection by WT and *spoh1*</u>

Seven-day-old *P. japonicum* seedlings were transferred to 0.8% water agar for starvation and placed next to 1-week-old Col-0 Arabidopsis seedlings, both grown vertically. The Arabidopsis main roots were carefully aligned with the *P. japonicum* roots, ensuring that both root tips were touching and oriented in the same direction. Haustoria numbers were counted after seven days of co-incubation.

<u>Haustorium xylem cell staining</u>

Root sections, including haustoria, were excised and immersed in 10% KOH, heated at 90 ºC for 15 min and washed 3 x with water, and were stained with 0.1% (w/v) Safranin-O (Wako Chemicals, Osaka, Japan) at 90 ºC for 5 min. Stained samples were then cleaned in a clearing solution containing 2.5 g/ml chloral hydrate (Nacalai Tesque) and 33% (v/v) glycerol for 15 min under vacuum. Haustorial structures were observed by light microscopy (Leica DMI3000 B).

Root architecture phenotyping

WT and *spoh1* seedlings were photographed after 27 days of incubation on agar or 2% sucrose. Total root length, main root length and lateral root number were quantified using Image J software (NIH).

Root extract and exudate preparation

WT and *spoh1* seedlings were incubated on agar or 4% sucrose for 2 weeks. Two-week-old Arabidopsis seedlings grown on 1/2 MS were used for root extract preparation. Roots were excised from the seedlings and ground in liquid nitrogen into fine powder, suspended in sterile water to make 5% (fresh w/v) stocks, and then gently mixed on a rotator mixer for 4 h to overnight at room temperature. The suspensions were then centrifuged at 20,400 x *g* for 10 min, and the supernatants were diluted with 0.8% agar or sterile water to the final working concentration for prehaustorium induction.

*P. japonicum* root exudates were collected by incubating WT or *spoh1* seedlings in water at 25 ºC under a 16 h light/8 h dark cycle with gentle shaking for 2 weeks. For sucrose-treated exudates, seedlings were incubated in 4% sucrose. The final concentration was standardized to 5 seedlings per ml. Arabidopsis root exudates were prepared by incubating 1-week-old seedlings in sterile water at 22 ºC under the same light condition with gentle shaking for 1 week. The final concentration was standardized to 2 seedlings per ml. For *P. japonicum* prehaustorium induction, 4 ml of exudate was applied to each well of a 12-well plate containing four *P. japonicum* 10-day-old seedlings, which have been starved for 3d, and incubated at 25 ºC under 16-h light/8-h dark cycles with gentle shaking for 1 week before observation.

*Striga* prehaustoria induction

*P. japonicum* root extracts (1% w/v final concentration) or root exudates were applied to the 96-well plate for *S. hermonthica* prehaustorium induction. Each well contained 100 µL of root extract or exudates and 50-100 *S. hermonthica* seedlings, which were treated with 10 nM strigol (or GR-24) for 24 h. The 96-well plates were then incubated at 25 ºC in the dark for 24 h before observation. The percentage of *S. hermonthica* seedlings forming prehaustoria in each well was calculated as the number of prehaustorium-forming seedlings divided by the number of germinated seedlings.

Whole-genome sequencing of *spoh1* populations

Eighty *spoh1* BC2F2 plants, each of which formed more than 3 spontaneous prehaustoria on 2% sucrose, were planted in pots. Leaves from these plants were harvested and pooled, and genomic DNA was extracted using the cetyltrimethylammonium bromide (CTAB) method (*26*). DNA quality was assessed using a bioanalyzer (Agilent), and sequencing was performed using an Illumina sequencer with 150 bp paired-end reads.

SNP identification for *spoh1*

SNP identification followed the published protocol (*27*) using CLC Genomic Workbench software (version 20). Briefly, sequence reads were quality filtered, removing those with a quality score lower than 0.05 and length shorter than 50 bp. Paired reads were mapped to the WT *P. japonicum* genome reference version 1 (*26*) with mismatch cost set to 2, length fraction at 0.6, and similarity fraction at 0.9. Background SNP filtering was applied by removing common low-stringency SNPs appearing in several other unrelated *P. japonicum* mutants with minimum coverage of 2, minimum count of 2, and minimum frequency of 10%. High-stringency SNPs were then called to identify target SNP(s) with a required variant probability of 100%, minimum frequency of 95%, minimum coverage of 5, and minimum count of 5. The resulting SNPs were annotated onto reference transcripts to identify nonsynonymous SNPs.

Plasmid construction

For overexpression constructs, the vector pUB-GW-GFP (*28*) was linearized by *Sal*I and *Xba*I, cleaving the ccdB region from the backbone. Linearized vector was purified using Wizard SV Gel and PCR Clean-Up System (Promega). The CDS of *PjGT72B1*, *PjWAS*, *Pjv1_00011317*, and *PjGT72B1* mutants and *AtGT72B1* were amplified from the cDNA of WT *P. japonicum* and Col-0 Arabidopsis cDNAs, respectively, using the primers listed in Table S2, with 15 bp overlaps at the end of the linearized pUB vector. The target genes were inserted using an In-Fusion HD Cloning Kit (Takara), and the resulting colonies were verified by Sanger sequencing. Empty vector was used as a control.

Golden Gate cloning technology was used for *PjGT72B1* promoter constructs. Lv1 constructs containing *35S::3xVenus-NLS* and mCherry-NLS with a *SmaI* restriction site at 7 bp upstream of the mCherry CDS were generated using Golden Gate modules (*29*). These two Lv1 fragments were assembled into the Lv2 construct pAGM4673 to create a backbone for promoter analysis. The *PjGT72B1* promoter region (3,866 bp) was amplified from genomic DNA using the primers listed in Table S2. The backbone was linearized by *Sma*I, and the *PjGT72B1* promoter region was inserted upstream of mCherry CDS by In-Fusion HD Cloning. The resulting colony was verified by Sanger sequencing.

Hairy root transformation and *spoh1* phenotype complementation

Hairy root transformation of *P.japonicum* was performed as described previously (*30*). Briefly, plasmid constructs were transferred into *Agrobacterium rhizogenes* strain LBA1334. Five-day-old seedlings were immersed in bacterial suspension (OD600 = 0.1) and sonicated for 20 sec. Sonicated seedlings were vacuum infiltrated for 7 min, then transferred to B5 agar medium containing 1% sucrose and 450 µM acetosyringone. Infected seedlings were incubated in the dark at 22 ºC for 2 days, then transferred to B5 agar medium containing 1% sucrose and 300 µg/ml cefotaxime, and incubated at 25 ºC under long-day conditions.

Arabidopsis mutant lines

pUB constructs containing *AtGT72B1*, *PjGT72B1* and mutated *PjGT72B1* were transferred into *Agrobacterium tumefaciens* strain AuL1. The *cs25067* mutant lines were transformed by floral dip method using these Agrobacterium strains. Positive transformants were identified by GFP florescence.

*spoh1* phenotype complementation

Hairy roots exhibiting GFP fluorescence were placed at the bottom of plastic petri dish, and covered with 0.8% agar containing 0.05 μM DMBQ, then vertically incubated for an additional 7 days before prehaustorium observation.

Generation of knock-out mutant roots by CRISPR/Cas9

PjGT72B1-KO and Pjv1_00011317-KO hairy roots were generated using the CRISPR/Cas9 system in WT *P. japonicum* following the Greifenhagen *et al.* protocol (*31*). Vectors containing Cas9 enzyme and AtU6P promoter were provided by Dr. Thomas Spallek. Guide RNAs for PjGT72B1 and Pjv1_00011317 were designed by CRISPOR (http://crispor.tefor.net/) (Table S2).

RT-qPCR analysis

Total RNA from individual hairy roots for complementation tests were extracted using a QIAGEN RNeasy Plant Mini kit, with genomic DNA elimination by DNase I (QIAGEN). First-strand cDNA was synthesized using the ReverTra Ace-a- kit (Toyobo; code FSK-101), and qPCR was performed using Thunderbird SYBR qPCR Mix kit (Toyobo; codeA4251K) on a Bio-Rad thermal-cycling system. A primer pair for PTB was used as an internal control in subsequent experiments. Primers for RT-qPCR are listed in Table S2.

Phylogenetic tree construction

UGTs in *P. japonicum* were identified by BLAST search using the conserved 44-amino acid PSPG box motif as a query sequence. Incomplete sequences were eliminated from the subsequent analysis. Arabidopsis UGT amino acid sequences were obtained from The Arabidopsis Cytochrome P450, Cytochrome $b_5$, P450 Reductase, ß-Glucosidase, and Glycosyltransferase Site (http://www.p450.kvl.dk/index.shtml). Full-length amino acid sequences were aligned by ClustalW, and Maximum Likelihood phylogenetic trees were constructed using MEGA-X software.

RNA-sequence analysis

Post-starvation 10-day-old WT and *spoh1* seedlings were incubated on 0.8% agar or 4% sucrose for 7 days. Root segments containing spontaneous prehaustoria of *spoh1* on sucrose were excised and used for subsequent RNA extraction. Similar parts of WT and *spoh1* roots were handled with the same procedure. Three replicates were conducted for each treatment. Total RNA was extracted using QIAGEN RNeasy Plant Mini kit with genomic DNA elimination by DNase I (QIAGEN). RNA quality was assessed with a 2100 bioanalyzer using an RNA 6000 Nano assay kit. Library construction and sequencing were performed by BGI using HiSeqX in paired-end mode with the a read length of 150bp. Reads were quality filtered and adaptor trimmed by Trimmomatic (ver. 0.39) (*32*) with the options of HEADCROP:15 LEADING:20 TRAILING:20 SLIDINGWINDOW:4:20 MINLEN:50 and fastqc (https://www.bioinformatics.babraham.ac.uk/projects/fastqc/), and mapped to the *P. japonicum* genome ver.1 (*26*) with the HiSAT2 program (*33*). Mapping results were converted to read counts by featureCounts in the R subread library, and differentially expressed genes were detected by edgeR. Self-organized mapping (SOM) clustering was performed as previously reported (*34*) using Kohonen (*35*). GO enrichment analysis was performed using ShinyGO (0.81) (*36*) using the top hit homolog from among Arabidopsis genes by a BLAST search of *P. japonicum* genes. Network analysis was performed by STRING (*37*) using Arabidopsis homolog genes.

<u>Confocal Microscopy</u>

For promoter analysis and protein subcellular localization, approximately 1 cm of hairy roots excisions exhibiting VENUS fluorescence were placed in glass-bottom Petri dishes (IWAKI, Japan) and covered with agar containing 0.4% Arabidopsis root extract. For mature haustoria induction, a Col-0 root was arranged carefully next to a hairy root under the agar layer and incubated at 25 ºC under long-day conditions until mature haustoria formed. 3xVENUS fluorescence was excited with a 514 nm laser, and the emission was detected between 525-560 nm using an Olympus FV3000 microscope. mCherry fluorescence was excited using a 543 nm laser, and the emission was detected between 570-640 nm. Hairy roots with a strong VENUS signal across the entire root were selected for these experiments.

<u>Purification of GST-PjGT72B1 expressed in *E. coli*</u>

The pDEST15 plasmid was used to generate a Glutathione-S-transferase (GST) fused protein in *E. coli* strain BL21. The pDEST15 backbone was linearized using *Not*I and *Sal*I, and the CDS of *PjGT72B1*, mutated *PjGT72B1* (H18Q, Y315F, H18Q and Y315F double mutant, D115A) as well as *AtGT72B1* were inserted using an In-Fusion Cloning Kit. Primers for gene amplification are listed in Table S2. The constructs were transformed into *E.coli* BL21 cells, which were grown at 37 ºC to an $OD_{600}$ of 0.6. Protein expression was induced with 0.1 mM IPTG at 20 °C for 24 h. To prepare the cell lysates, pellets from 50 mL of culture were resuspended into 1 ml of extraction buffer (140 mM NaCl, 10 mM $Na_2HPO_4$, 1.8 mM $KH_2PO_4$, pH 7.5), sonicated for 5 seconds, followed by a 5-second pause on ice, and the sonication cycle was repeated 10 x per sample. Cell lysates were then centrifuged at 11,000 x *g* for 10 min at 4ºC. Supernatants were then loaded onto a Glutathione-Superflow Resin column (Takara) to purify GST-fused proteins. Elution was performed with a glutathione-containing buffer (33 mM glutathione in 50 mM Tris-HCl, pH 7.0). The eluted protein was desalted using a PD MiniTrap G-25 column (Cytiva). Protein concentrations were measured using the Bio-Rad Protein assay, using Immunoglobulin G to generate a standard curve.

GST-fused GT72B1 *in vitro* enzyme assay

*In vitro* reaction mixture (200 µl) contained 2 µg of recombinant protein, 14 mM 2-mercaptoethanol, 5 mM UDP–glucose and 1 mM substrate. Enzyme assays for OGT activity were carried out in a buffer containing 100 mM Tris-HCl (pH 7.0). For NGT activity, the reaction buffer contained 100 mM Tris-HCl (pH 8.0). Both assays were carried out at 30°C for 1 h before analysing the glucosylated product peak by Reverse phase HPLC (Alliance Waters e2695) using a Columbus 5 µm C18 column. HPLC mobile phase was 0.1% trifluoroacetic acid (TCA) in water for 2 min, followed by 28 min linear gradient of acetonitrile with 0.1% TCA from 0 to 50%, then 95% acetonitrile for 5min, followed by 100% water for 5 min at 0.8 ml/min to separate the glucose conjugates from aglycones. OGT and NGT product peaks were detected by UV absorbance at 296 and 300 nm, respectively. The enzyme assay products were analyzed by LC-MS. Chromatographic separations were conducted on UHPLC "Vanquish Duo" (Thermo Fisher Scientific, San Jose, USA) with 3 µL injection, equipped with a InertSustain C18 column (100 by 2.1 mm i.d. 3 µ particle size, GL Science, Japan). The mobile phase consisted of solvent A (0.1% formic acid in water) and B (0.1% formic acid in acetonitrile). Following gradient was applied at a flow rate of 200 µL min-1: 0-1 min, 100%; 1-2 min, from 100% to 90% A; 2-16 min, from 90% to 60% A; 16-22 min, from 60% to 0% A; 22-25 min, to 0% A, column wash; 25-30 min, to 100% A for equilibration of the column. Compounds were detected from m/z 150-

600 by L LTQ XL™ Linear Ion Trap Mass Spectrometer (Thermo Fisher Scientific, San Jose, USA) using full scan mode covering positive (sid = 20) and negative (sid = 5; 30) ion detection. The chromatograms were analyzed by Xcalibur software version 4.1 (Thermo Fisher Scientific, San Jose, USA), the m/z value, retention time and detection mode information of target peaks were extracted.

<u>Synthesis of glucosylated HIFs</u>

Vanillin 4-*O*-β-D-glucopyranoside (vanillin-Glc) was synthesized as previously described (*38*). Syringic acid 4-*O*-β-D-glucopyranoside (SyA-Glc), ferulic acid 4-*O*-β-D-glucopyranoside (Fer-Glc), and sinapic acid 4-*O*-β-D-glucopyranoside (SinA-Glc) (fig. S1) were synthesized via the Koenig-Knorr glycosylation of methyl synringate (methyl 4-hydroxy 3,5-dimethoxybenzoate), ethyl ferulate (ethyl 4-hydroxy-3-methoxycinnamate), and ethyl sinapate (ethyl 4-hydroxy-3,5-dimethoxycinnamate), respectively, followed by alkaline deprotection. Briefly, the protected aglycone (methyl syringate, ethyl ferulate, or ethyl sinapate, 1.0 equiv.) was mixed with 2,3,4,6-tetra-*O*-acetyl-α-D-glucopyranosyl bromide (1.0 equivalent) in quinoline at 0 °C. Silver oxide (1.0 equiv.) was then added under vigorous stirring. The reaction mixture was stirred at ambient temperature for 1.5 h, filtered through a Celite pad, and extracted with ethyl acetate. The extract was sequentially washed with 1 M hydrochloric acid, saturated aqueous sodium bicarbonate, and brine, dried over anhydrous sodium sulfate, and concentrated under reduced pressure to yield the crude protected glucosides. The glucosides were then deprotected in a 1:2 (v/v) mixture of ethanol and 0.5 N aqueous sodium hydroxide solution (for SyA-Glc) or in a sodium methoxide solution (approximately 5%) in methanol (for Fer-Glc and SinA-Glc) at 0˚C. After neutralization with acetic acid, the reaction mixture was concentrated under reduced pressure and purified by flash chromatography using a methanol/dichloromethane eluent system to yield the pure glucosides. The purity and structural identity of the glucosides were confirmed by standard NMR and LC-MS. SyA-Glc (pale yellow crystals): $^1$H NMR (400 MHz, DMSO-$d_6$) δ 7.19 (s, 2H, H-2, H6), 5.1 (d, *J* = 6.9 Hz, 1H, H-1′), 3.78 (s, 6H, 2 × $OCH_3$), 3.53 (dd, *J* = 5.0, 4.6 Hz, 1H, H-6′a), 3.43 (dd, *J* = 6.0, 6.0 Hz, 1H, H-6′b), 3.41–3.20 (m, 3H, H-2′, H-3′, H-5′), 3.17 (m, 1H, H-4′). LC-MS (ESI) calculated for $C_{15}H_{20}O_{10}$ + $CH_3COO^-$ [$(M+CH_3COO)^-$]: 419; found: 419. FerA-Glc (colorless crystals): $^1$H NMR (400 MHz, DMSO-$d_6$) δ 7.59 (d, *J* = 15.9 Hz, 1H, H-α), 7.37 (d, *J* = 1.8 Hz, 1H, H-2), 7.22 (dd, *J* = 1.8, 1.4 Hz, 1H, H-6), 7.08 (d, *J* = 8.7 Hz, 1H, H-5), 6.58 (d, *J* = 16 Hz,

1H, H-β), 4.97 (d, $J$ = 7.3 Hz, 1H, H-1′), 3.80 (s, 3H, $OCH_3$), 3.65 (d, $J$ = 11.4 Hz, 1H, H-6′a), 3.38 (m, 1H, H-5′), 3.22–3.27 (m, 2H, H-2′, H-3′), 3.15 (m, 1H, H-4′), 3.42 (dd, $J$ = 11 Hz, 1H, H-6′b). $^1$H NMR (400 MHz, DMSO-$d_6$) δ 7.55 (d, $J$ = 16 Hz, 1H, H-α), 7.03 (s, 2H, H-2, H6), 6.62 (d, $J$ = 15.6 Hz, 1H, H-β), 5.05 (s, 1H, H-1′), 3.75 (s, 6H, 2 × $OCH_3$), 3.53 (d, $J$ = 9.1 Hz, 1H, H-6′a), 3.48–3.18 (m, 5H, H-2′, H-3′, H-5′, H-6′b), 3.15 (m, 1H, H-4′). LC-MS (ESI) calculated for $C_{17}H_{22}O_{10}$ + $CH_3COO^-$ [$(M+CH_3COO)^-$]: 445; found: 445.

LC-MS analysis of root exudates

WT or *spoh1* were grown in 50 ml water or 4% sucrose for 14 days at 25 ˚C with gentle shaking at a density of 5 seeds per ml. Root exudates were collected as above and freeze dried. Targeted phenolic analysis was performed as described previously (*39*). A Waters Acquity UPLC I-Class System (Waters, Milford, MA, USA) equipped with a Binary Solvent Manager and Sample Manager was employed as a chromatographic system coupled to a Xevo TQ-S tandem quadrupole mass spectrometer (Waters MS Technologies, Manchester, UK) with electrospray (ESI) ionization interface. Five µL of root exudates (semi-polar and polar franctions) were injected and separated on an Acquity UPLC BEH C18 column (2.1 3 100 mm, 1.7 mm particle size, Waters, Milford, MA, USA) with 15 mM formic acid in both water (A) and acetonitrile (B). At a flow rate of 300 µL per min and a column temperature of 40 ˚C, a gradient of 0 min, 5% B; 2 min, 5% B; 32 min, 18% B; 60 min, 24% B; 65 min, 100% B was used as mobile phase. Compounds were measured in the ESI ion source of the tandem mass analyzer operating as described previously (*40*). Mass data of phenolic compounds were acquired in multiple reaction monitoring (MRM) mode. The MassLynx software, ver 4.1 (Waters), was used to control the instrument as well as to acquire and process MS data.

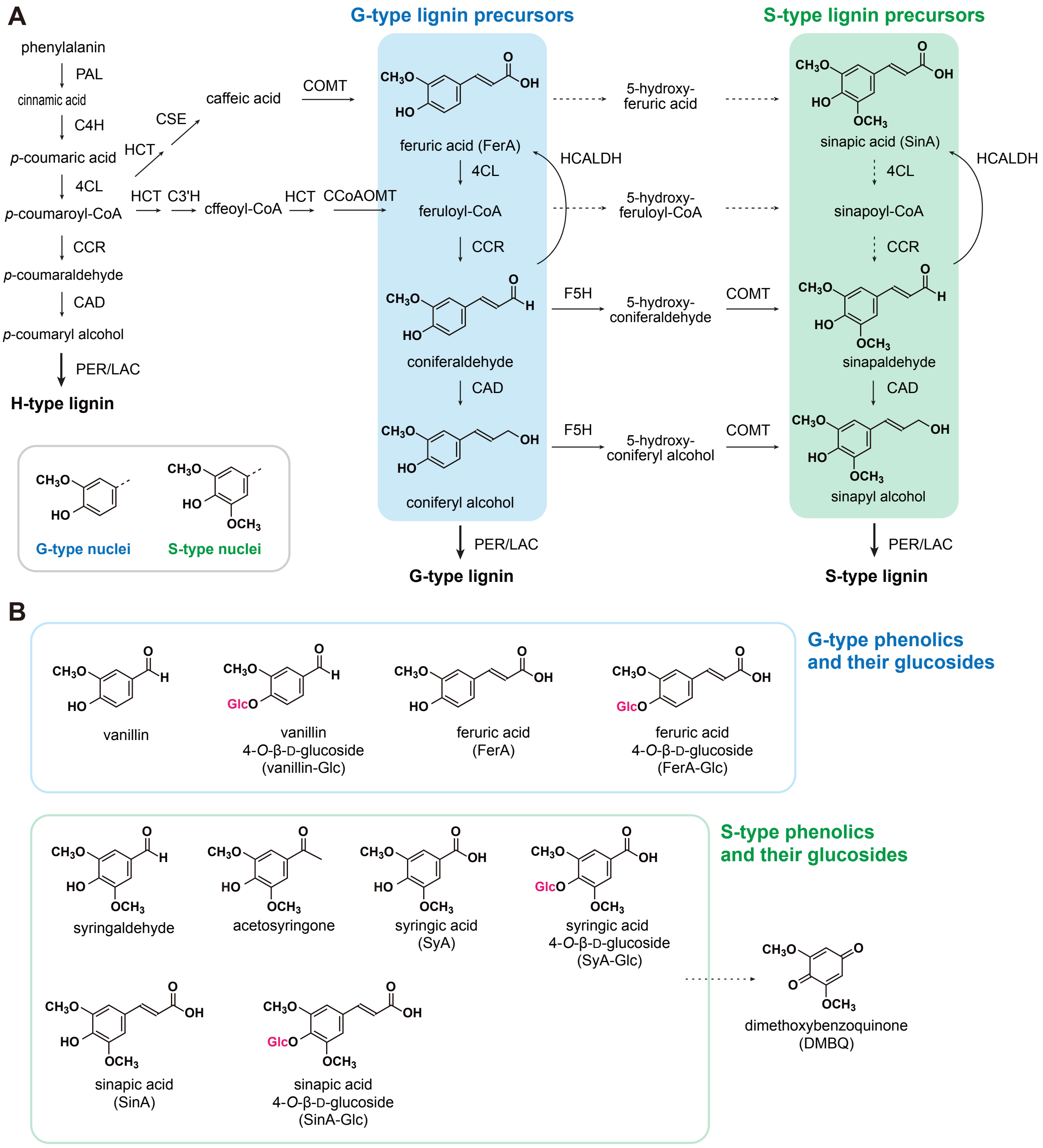


**fig. S1. Lignin biosynthesis pathway and chemical structures of HIFs and their glucosides.** **(A)** The pathway of G- and S-type monolignol biosynthesis. **(B)** The structures of known HIFs and their glucosides.

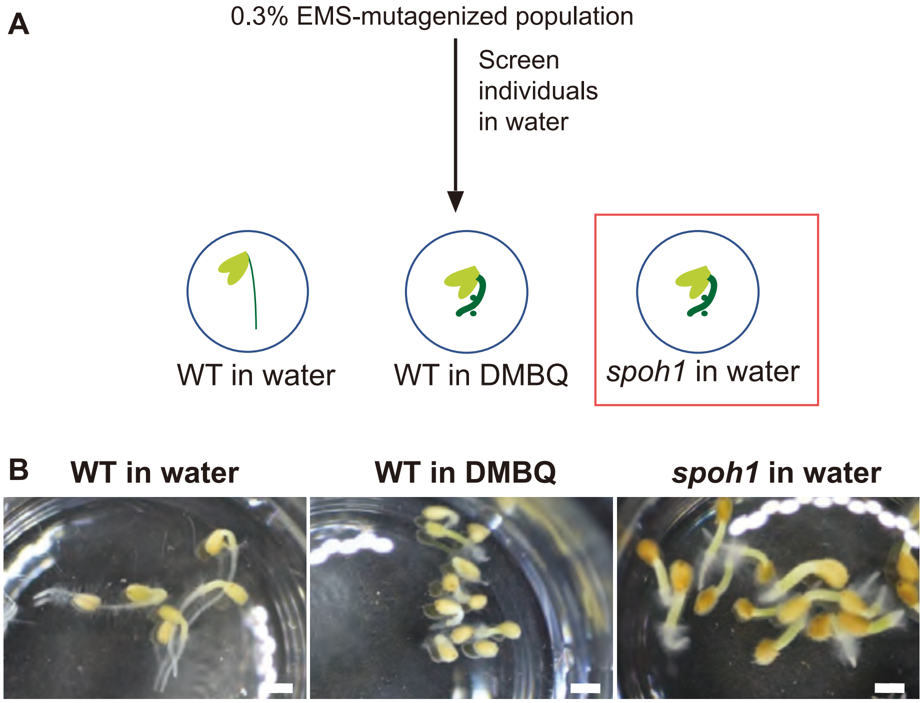


**fig. S2. Isolation of *spoh1* mutant.**

**(A)** Schematic representation of the screening procedure for isolating *spontaneous prehaustorium 1* (*spoh1*) mutant. An EMS-mutagenized population of *P. japonicum* was incubated in water, and mutants showing a phenotype similar to DMBQ-treated samples were selected. **(B)** Representative images of wild type (WT) seedlings in water or DMBQ and *spoh1* seedlings in water. Scale bars = 1 mm.

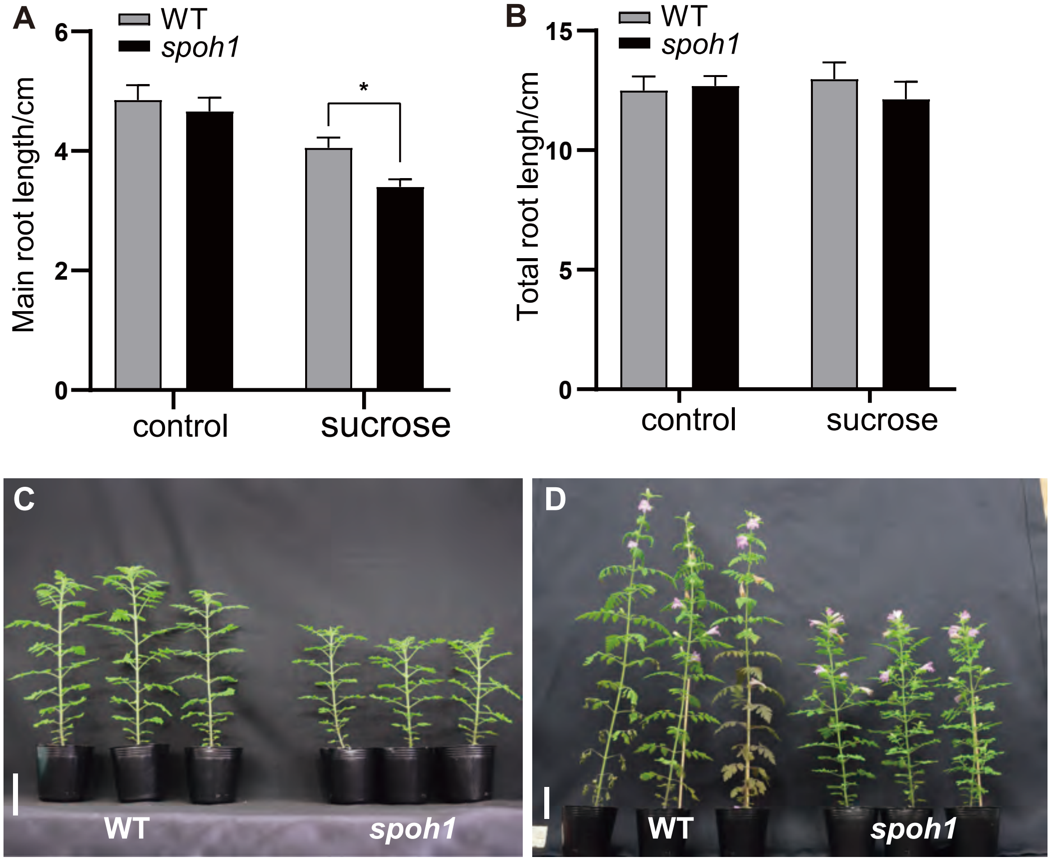


**fig. S3.** ***spoh1* phenotype**

**(A)** Main root length of WT and *spoh1* seedlings grown on media with (sucrose) or without (agar) 2% sucrose for 2 weeks. **(B)** Total root length of WT and *spoh1* under the same condition as (A). Data are mean ± SE from three experiments; n = 30. Student's t test, unpaired, two-tailed (*P < 0.05). **(C, D)** Representative phenotypes of WT and *spoh1* plants during vegetative growth (45-day-old) and reproductive growth (75-day-old). Scale bar: 5 cm.

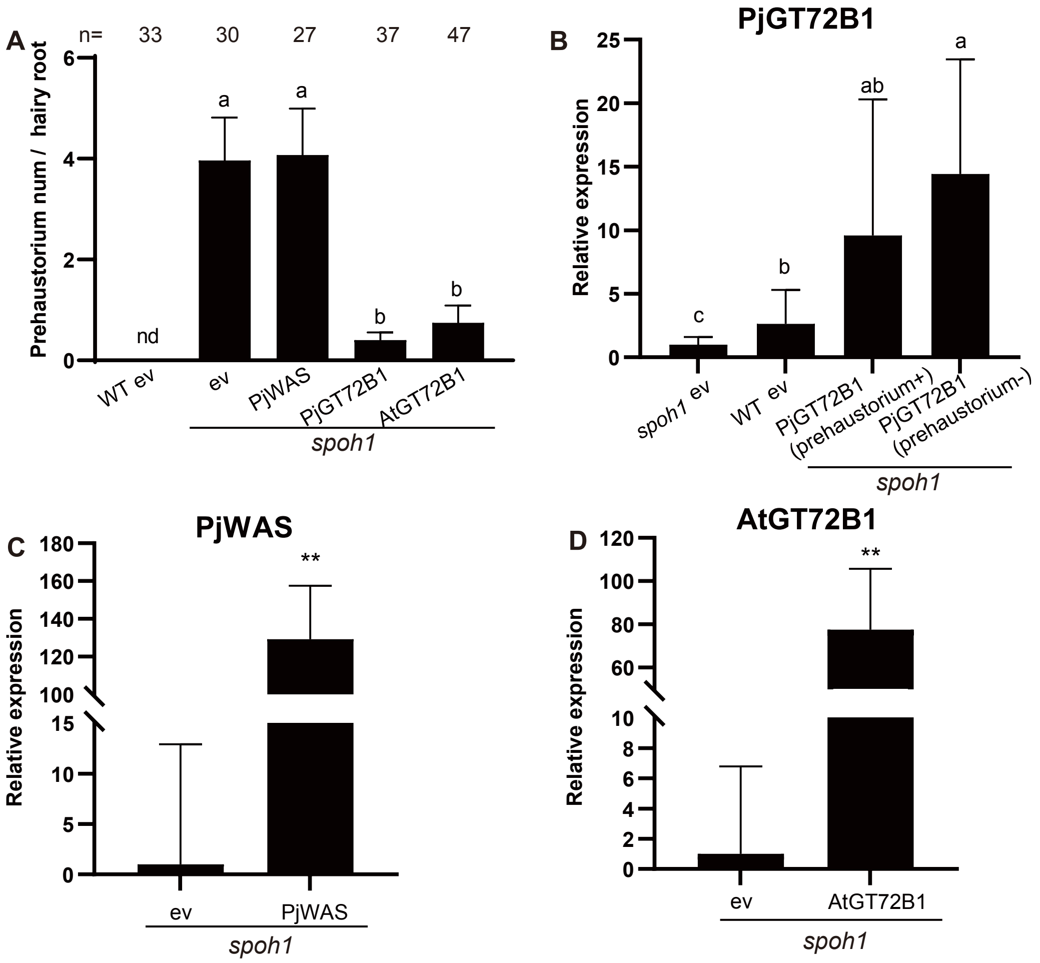


**fig. S4. *spoh1* complementation test.**

**(A)** Number of prehaustoria formed on hairy roots in complementation test after 7 days of induction with 0.05 μM DMBQ. Hairy roots with strong GFP signals were selected to minimize chimeric affects. Sample size (n) is indicated at the top of each bar in the chart. **(B-D)** Relative expression levels of Pj*GT72B1* in Pj*GT72B1* overexpressing roots forming prehaustoria (prehaustorium+) or not (prehaustorium-) (B), Pj*WAS* in Pj*WAS*-overexpressing roots (C), and At*GT72B1* in At*GT72B1*-overexpressing roots (D), as determined by qRT-PCR. Data are mean ± SE from three representative experiments. Significant difference between samples were calculated with One-way ANOVA (A, B) or student's t test, unpaired, two-tailed (**$P < 0.01$) (C, D). Hairy roots transferred with an empty vector (ev) served as controls. Data are mean ± SE from three representative experiments.

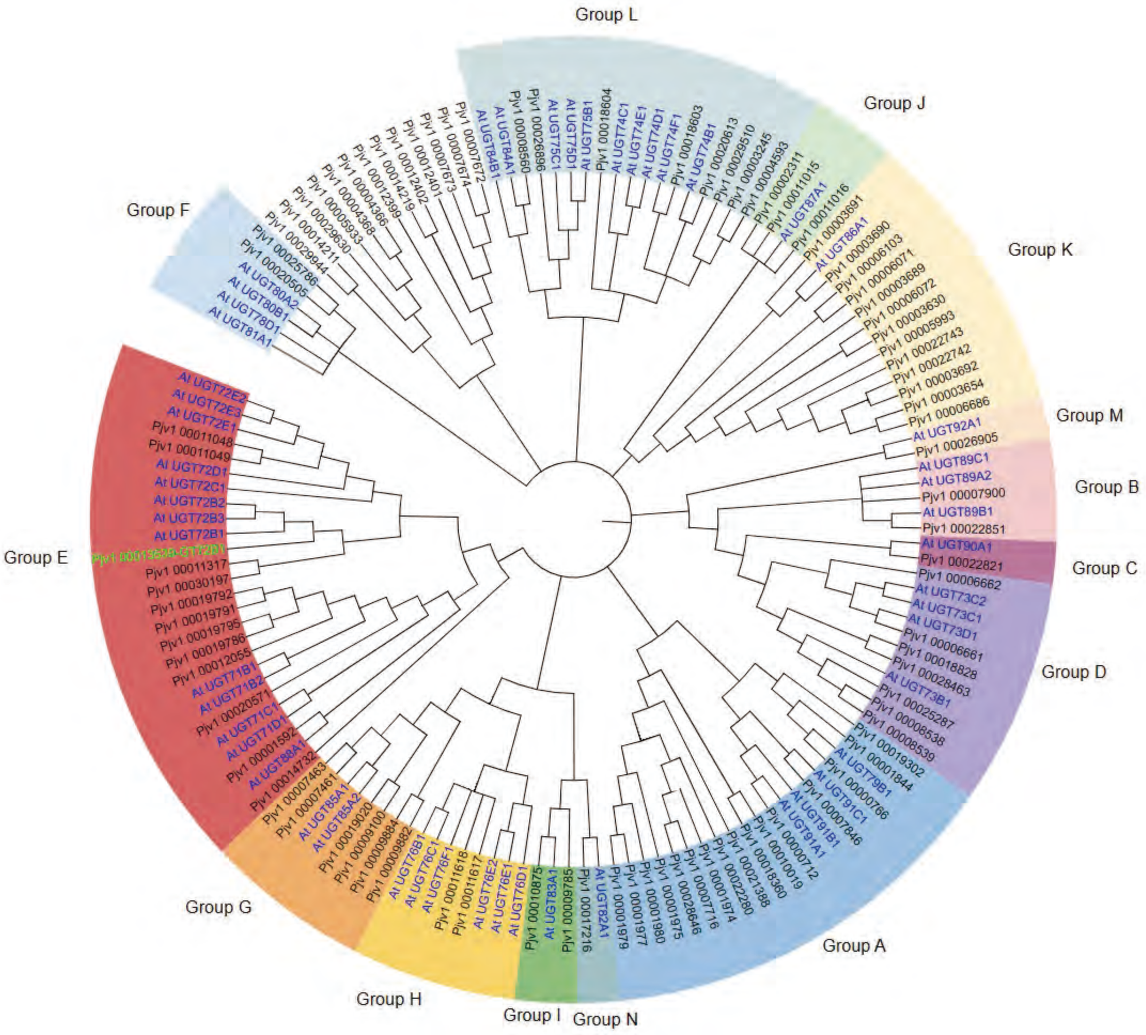


**fig. S5. Phylogenetic tree of UGTs from *P. japonicum* and *A. thaliana***

The phylogenetic tree was constructed using 52 Arabidopsis UGTs (blue) and 90 *P. japonicum* UGTs (grey). The *spoh1* gene is highlighted in green.

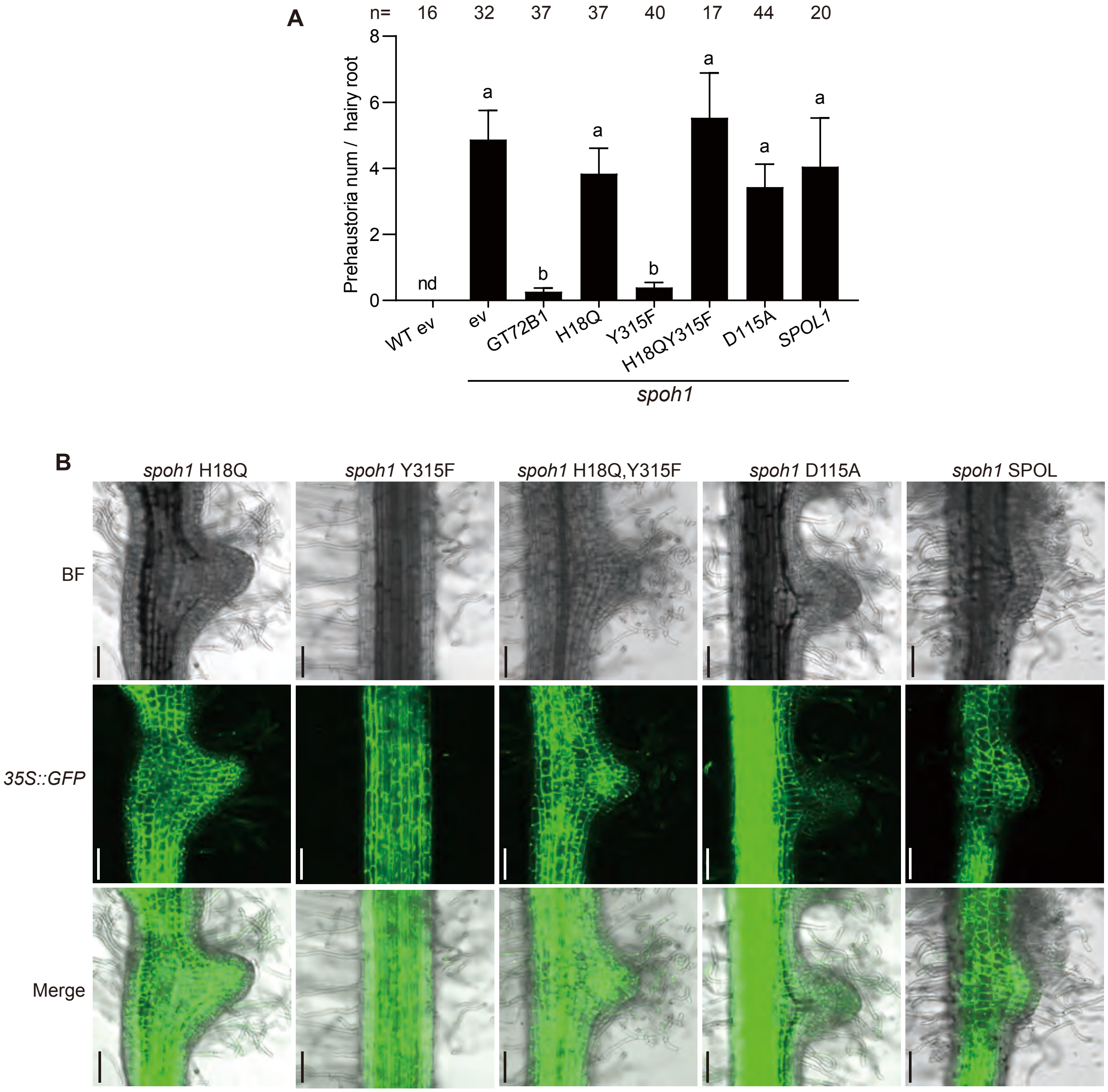


**fig. S6. Phenotype of *spoh1* hairy roots overexpressing various *PjGT72B1* mutant constructs.**

(A) Prehaustorium number in transgenic hairy roots after 7-day incubation on 0.05 μM DMBQ medium. Hairy roots with strong GFP signals were selected to minimize chimeric affects. ev represents the empty vector control. Data are mean ± SE from three representative experiments. The sample size (n) is indicated at the top of each bar. Significant difference was detected by One-way ANOVA. (B) Phenotypes of hairy roots overexpressing Pj*GT72B1* mutants or *SPOL1* in the *spoh1* background. Hairy roots were treated with 0.05 μM DMBQ for 1 week. Scale bar: 100 μm.

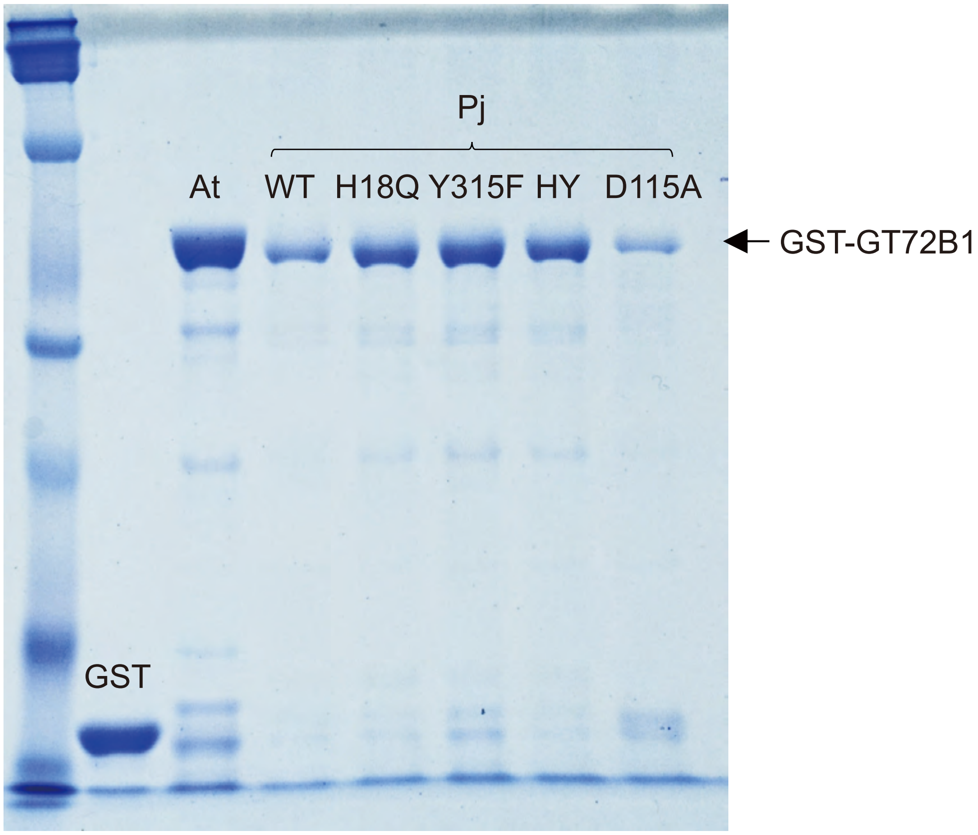


**fig. S7. Purification of GST-fused GT72B1 enzymes.**
GST-fused enzymes were expressed in *E.coli* and purified using a GST-affinity column. The purified proteins were subjected to SDS-PAGE and visualized with Coomassie Brilliant Blue staining. Black arrows indicate the bands corresponding to the GST-fused proteins. HY represents the H18Q and Y315F double mutant.

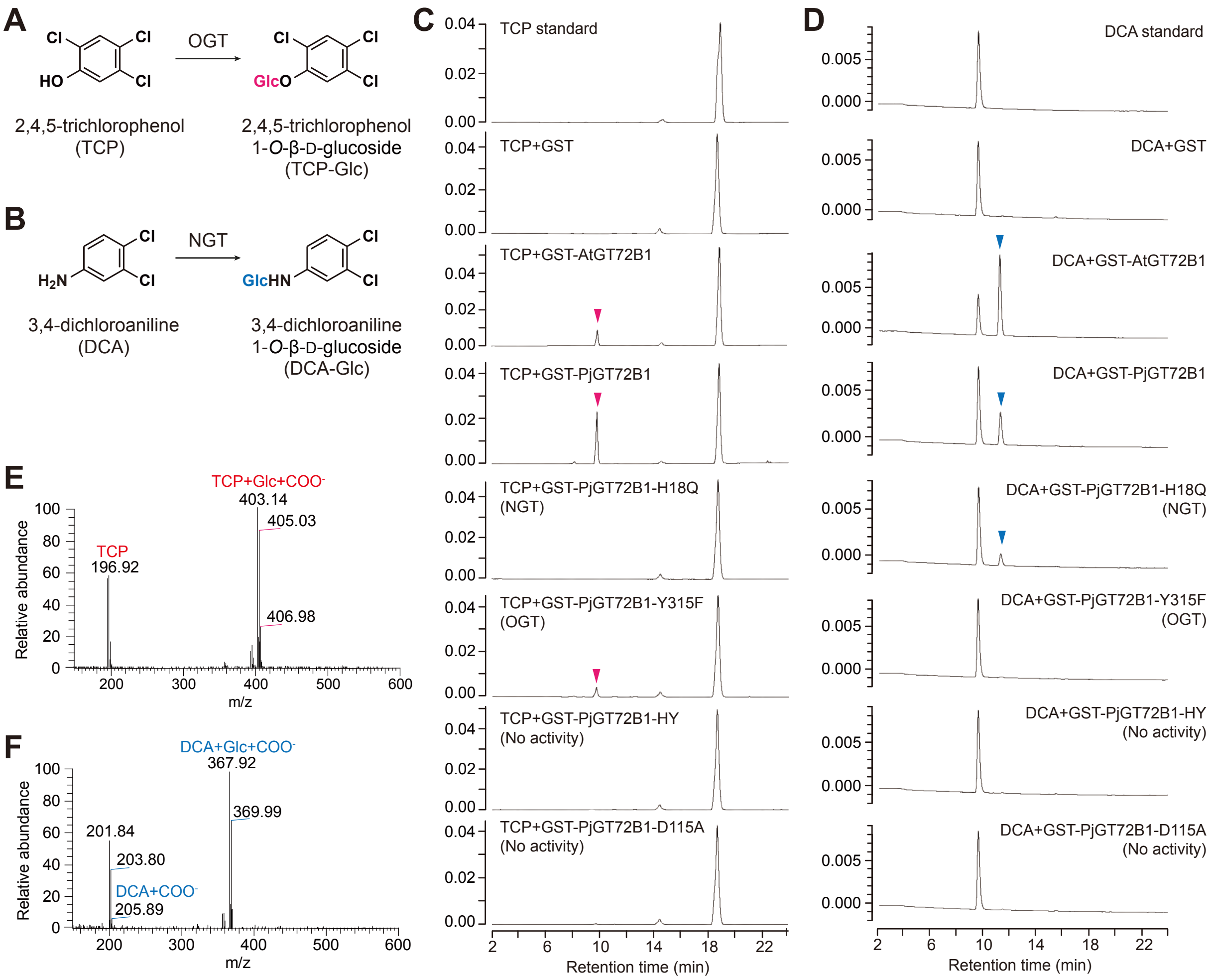


**fig. S8. NGT and OGT activities of purified PjGT72B1 proteins.**

(A, B) Chemical structures of TCP (A) and DCA (B), the substrates used for enzyme assays to determine OGT and NGT activities, respectively. (C, D) HPLC chromatograms of enzyme reaction products using TCP (C) and DCA (D) as substrates. Product peaks are indicated by red arrowheads. (E, F). MS spectra of the product peaks for TCP (E) and DCA (F) confirmed by LC-MS analysis.

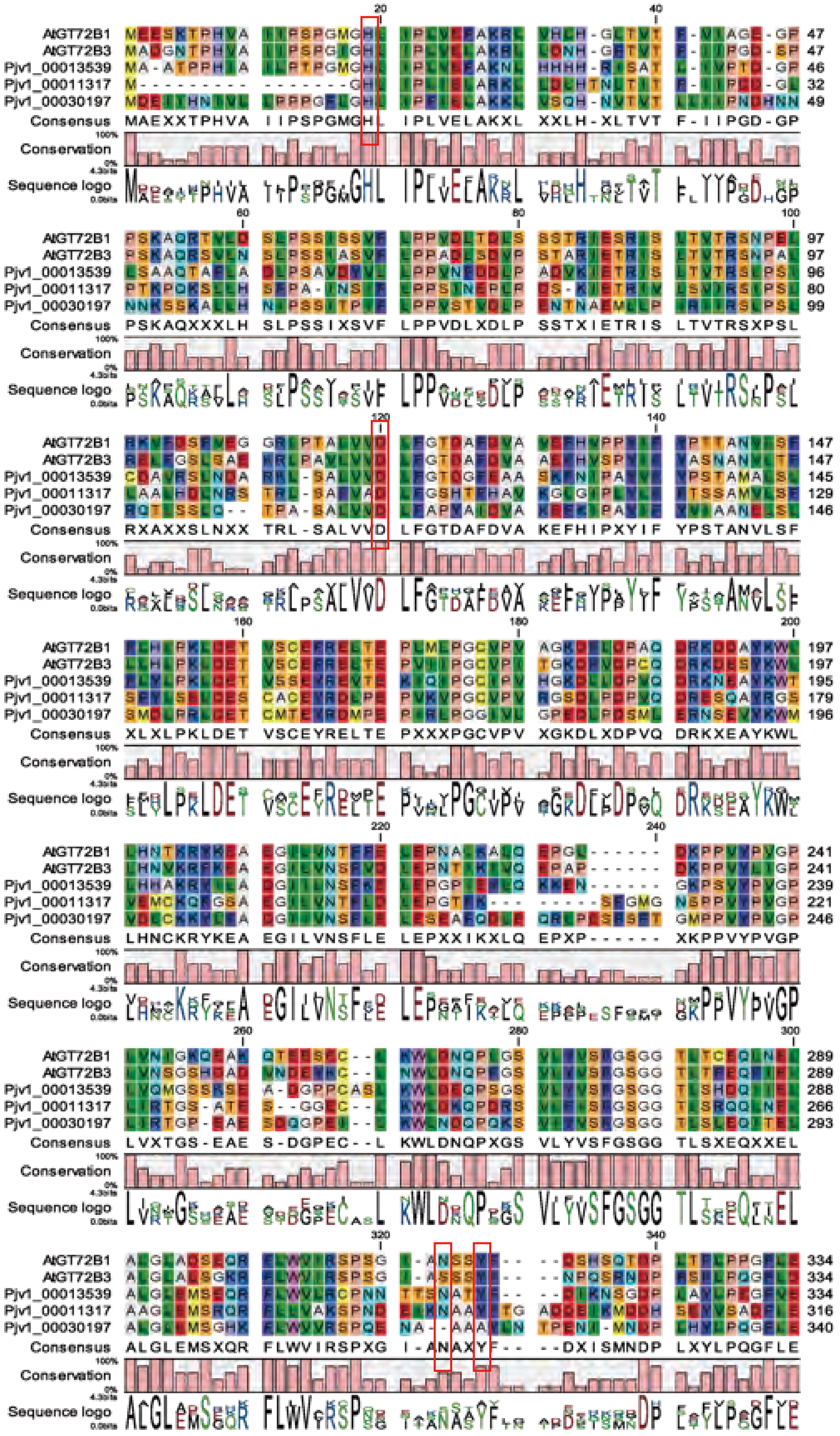

AtGT72B1
AtGT72B3
Pjv1_00013539
Pjv1_00011317
Pjv1_00030197
Consensus
Conservation
Sequence logo
MAEXXTPHVA IIPSPGMGHL IPLVELAKXL XXLH-XLTVT F-IIPGD-GP
PSKAQXXXLH SLPSSIXSVF LPPVDLXDLP SSTXIETRIS LTVTRSXPSL
RXAXXSLNXX TRL-SALVVD LFGTDAFDVA KEFHIPXYIF YPSTANVLSF
XLXLPKLDET VSCEYRELTE PXXXPGCVPV XGKDLXDPVQ DRKXEAYKWL
LHNCKRYKEA EGILVNSFLE LEPXXIKXLQ EPXP------ XKPPVYPVGP
LVXTGS-EAE S-DGPEC--L KWLDNQPXGS VLYVSFGSGG TLSXEQXXEL
ALGLEMSXQR FLWVIRSPXG I-ANAXYF-- --DXISMNDP LXYLPQGFLE

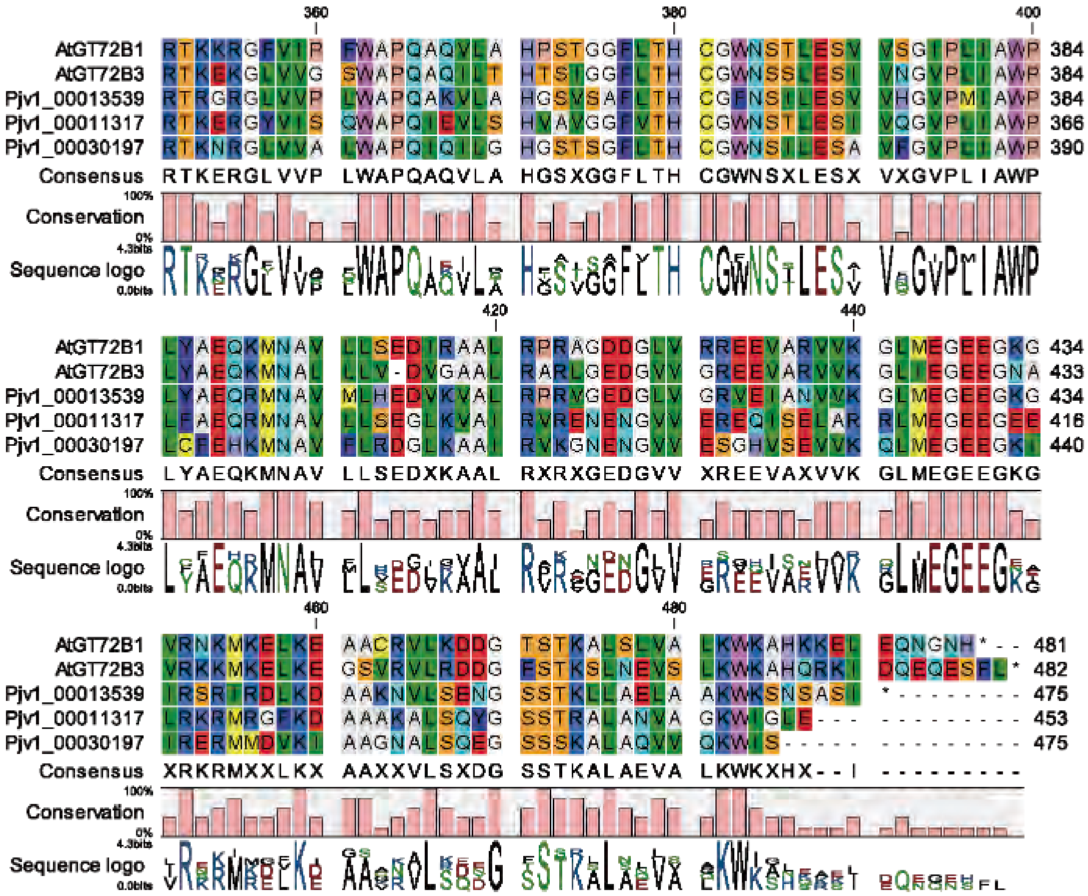

**fig. S9. Alignment of GT72B amino acid sequences.**
Amino acid sequences of AtGT72B1 (bifunction), AtGT72B3 (OGT), Pjv1_00013539 (SPOH1), Pjv1_00011317 (SPOL1) and Pjv1_00030197 (SPOL2) were aligned. Key residues, His19, Asp117, Asn312 and Tyr315 are highlighted with red rectangle.

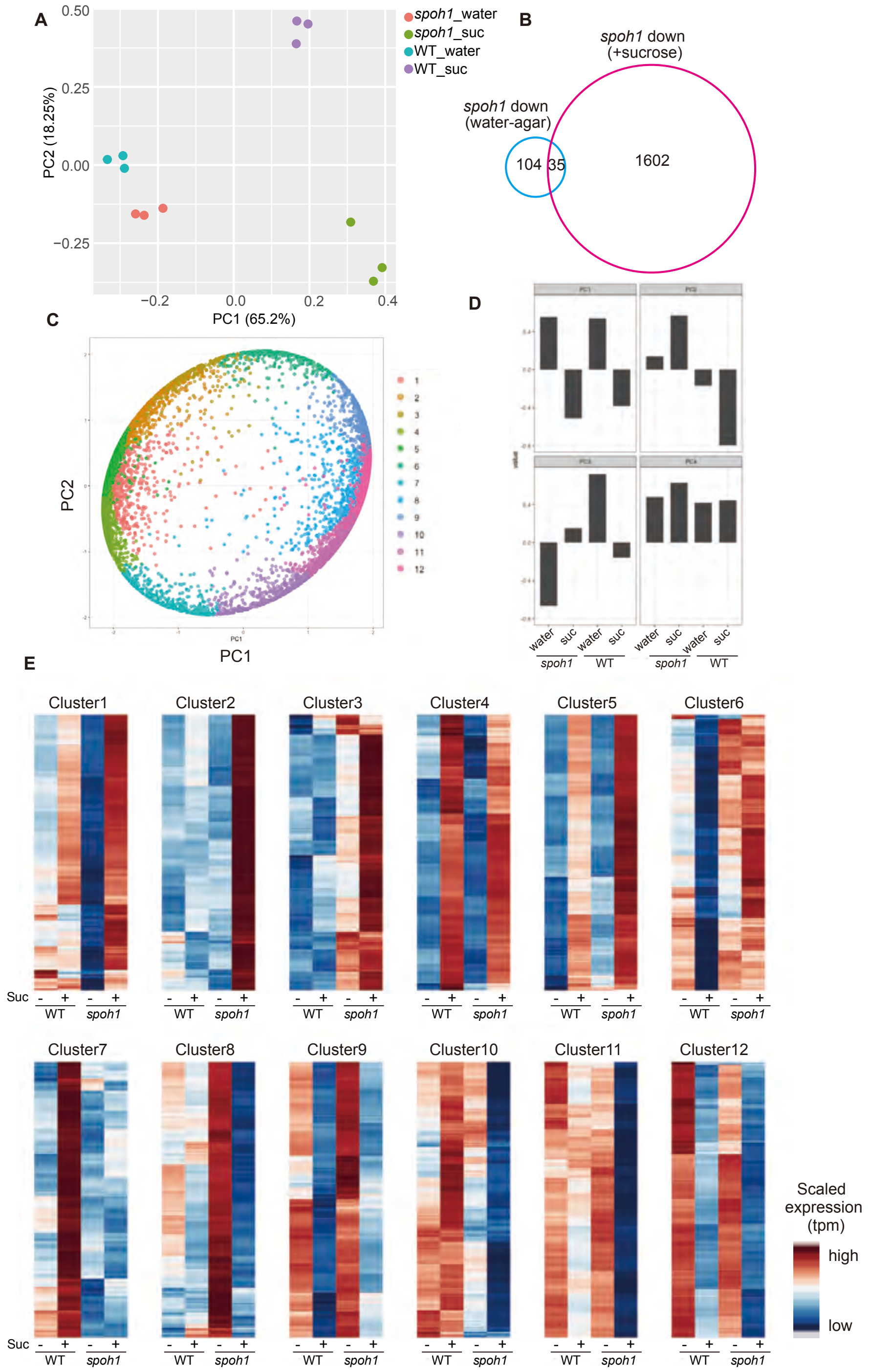


**fig. S10. RNA-sequencing of *spoh1* and WT with or without sucrose**

**(A)** Principal component (PC) analysis of RNA sequencing results. (B) Venn diagram showing the down-regulated genes in *spoh1*. (C) SOM clustering analysis of 6968 DEGs. (D) Loading of each PC used for the SOM clustering. (E) Heatmaps for each cluster after SOM clustering (Cluster 3 heatmap is the same as shown in Fig. 4D).

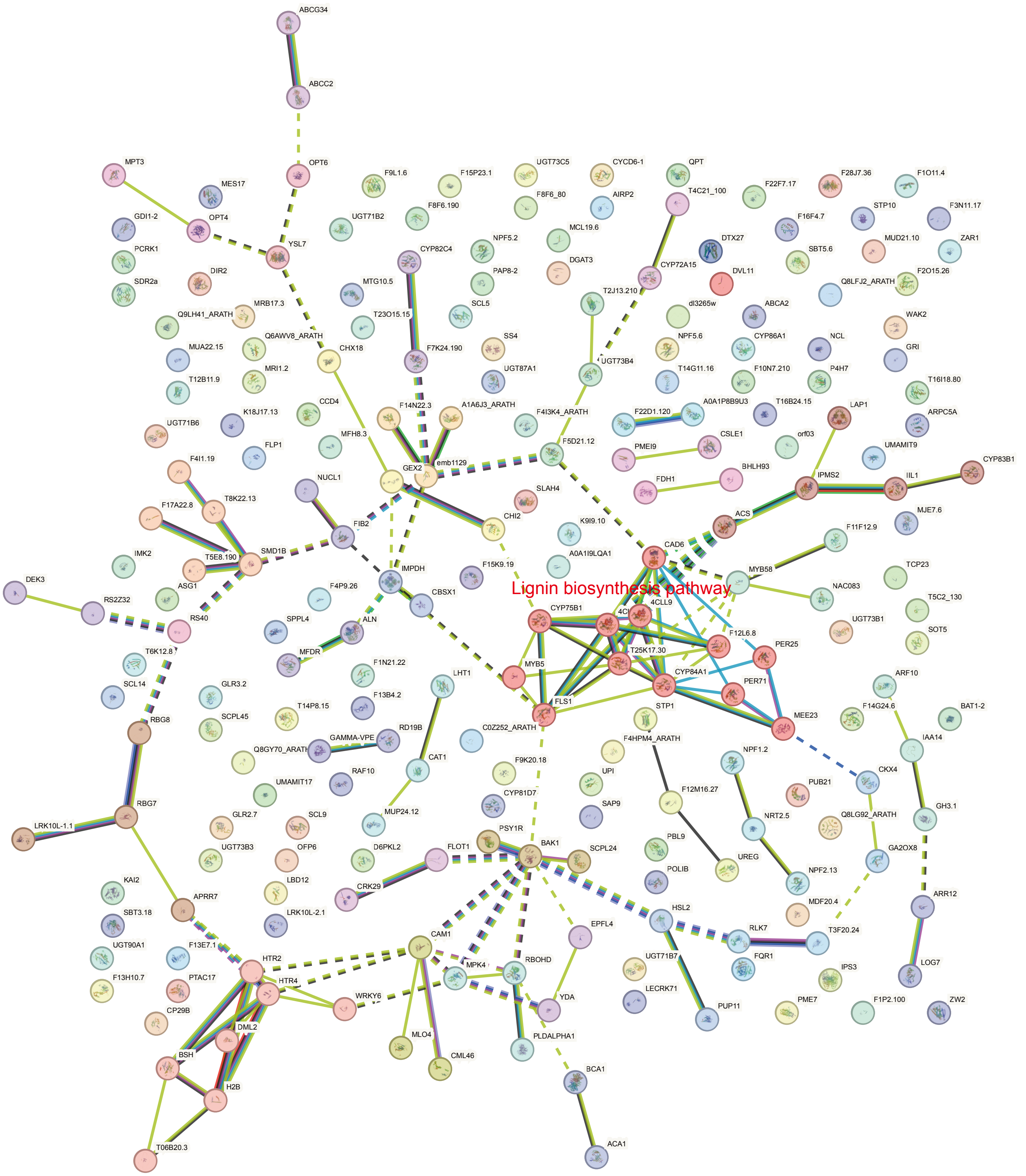


**fig. S11. STRING network analysis of genes involved in Cluster 3.**

The closest Arabidopsis homologs of cluster 3 genes were subjected to STRING network analysis. Lignin-related genes are highlighted in red.

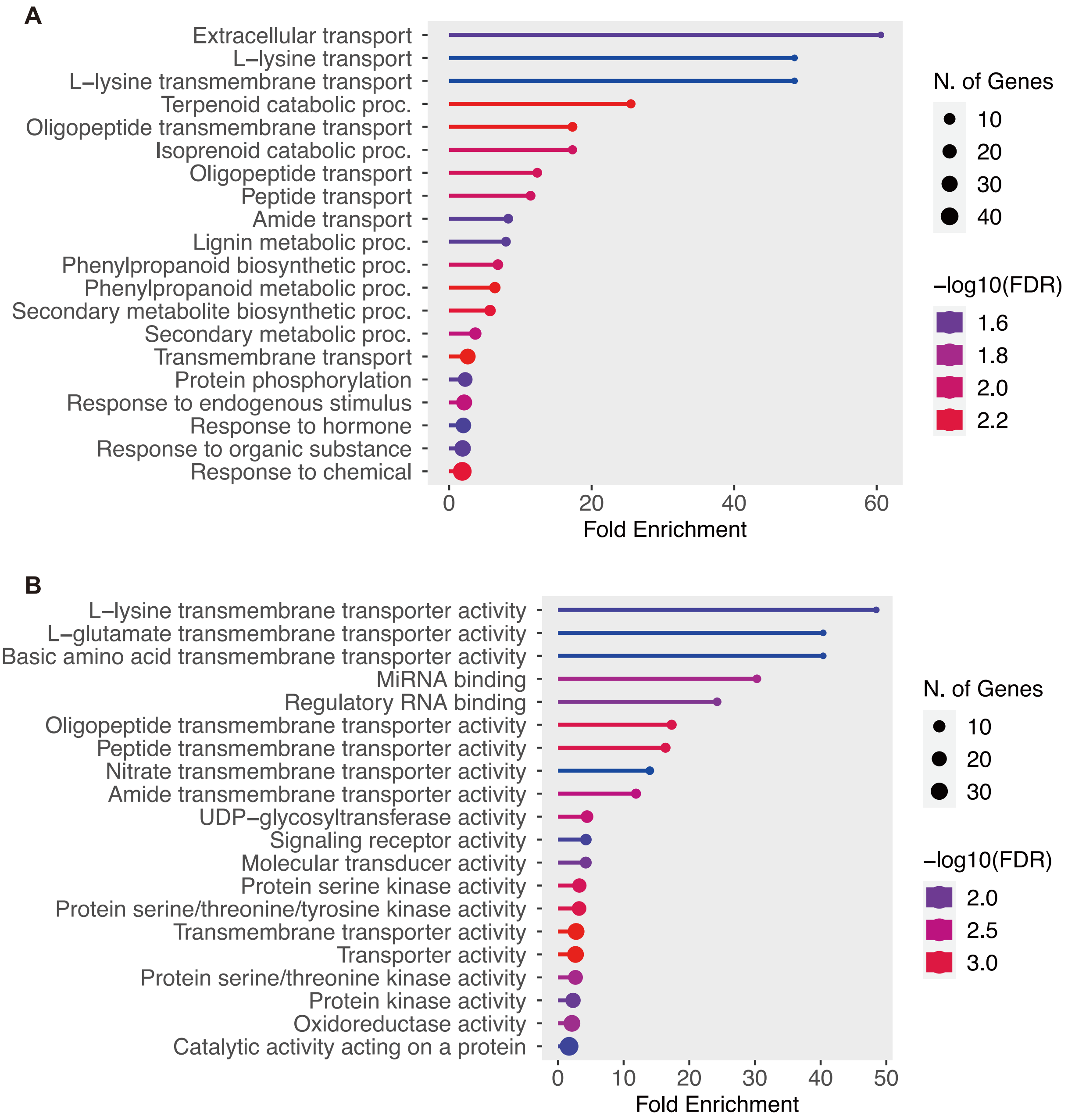


**fig. S12 GO enrichment analysis of cluster 3 genes**
GO enrichment analysis results for Biological Processes. B. GO enrichment analysis results for Molecular Functions.

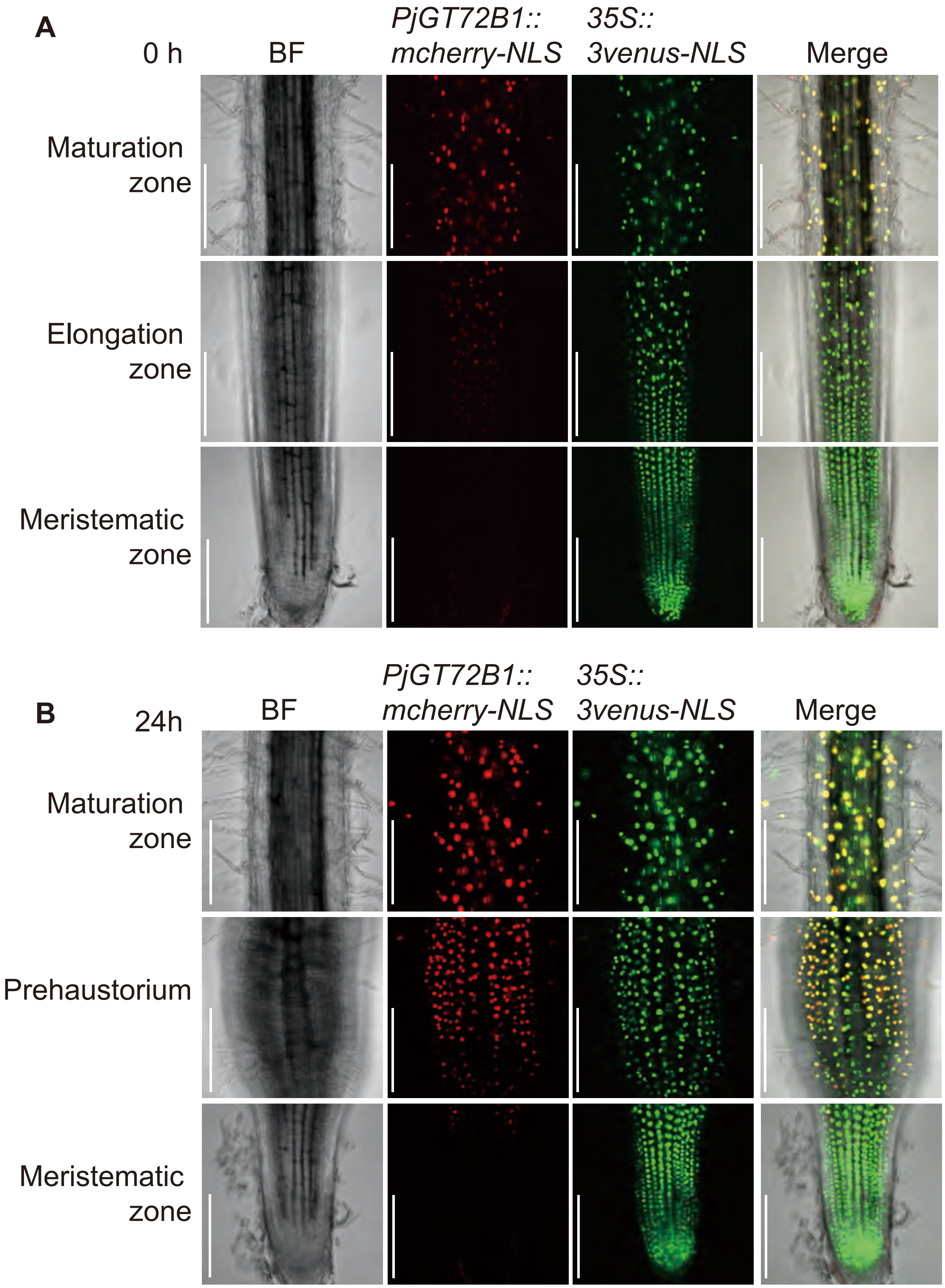


**fig. S13. Expression patterns of *PjGT72B1* in *P. japonicum* roots.**

**(A, B)** *PjGT72B1* expression pattern in different root tissues at 0h (A) and 24h (B) after prehaustorium induction. The mCherry panel indicates *PjGT72B1* expression, while the Venus panel shows marker gene expression. Hairy roots were treated with 0.4% Arabidopsis root extract. Scale bar: 100 µm.

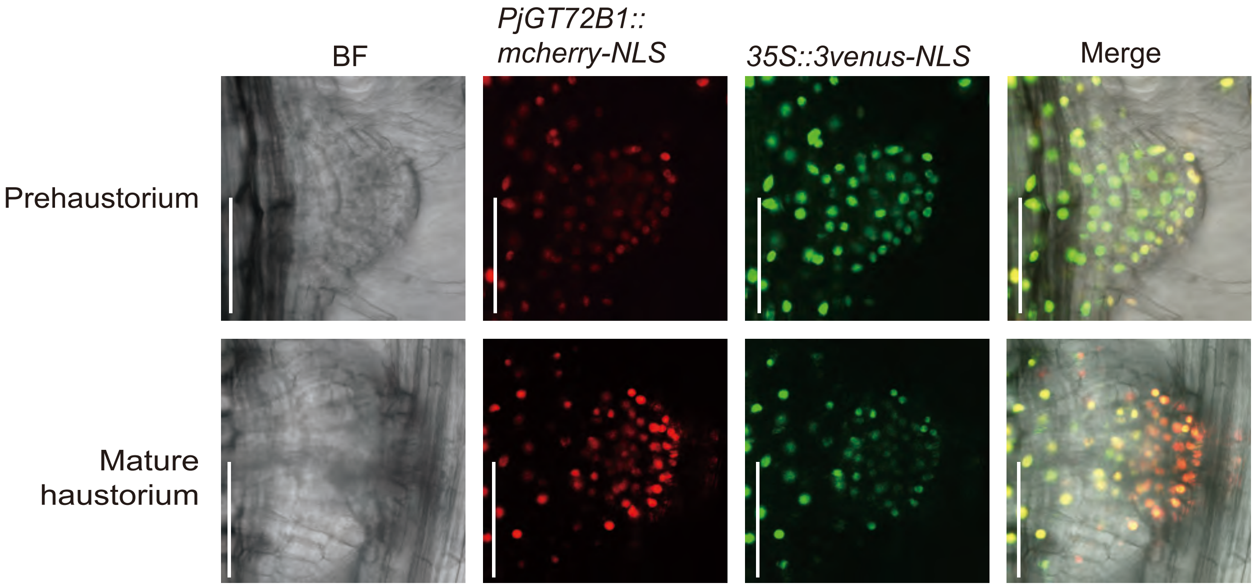


**fig. S14. Expression patterns of *PjGT72B1* in a prehaustorium and a mature haustorium.** The PjGT72B1::mCherry-NLS construct was introduced into *P. japonicum* hairy roots, which were then used to infect Arabidopsis roots. mCherry fluorescence was observed in a prehaustorium (prior to host invasion), and a mature haustorium (after host invasion). Venus-NLS was used as a transformation marker. Scale bar: 100 µm.

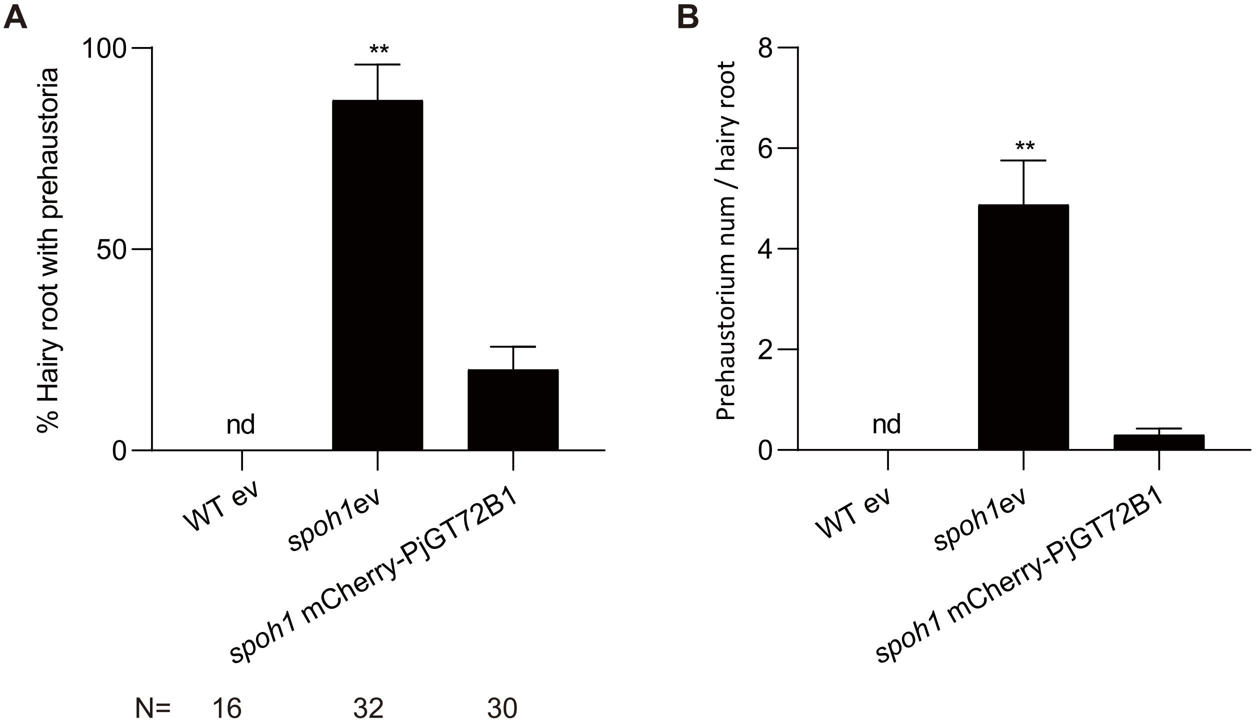


**fig. S15. mCherry fused PjGT72B1 complements *spoh1* phenotype**.
(A, B) *spoh1* hairy roots transformed with empty vector (ev) or mCherry-fused PjGT72B1 were incubated in 0.05 μM DMBQ medium for 7 days. Percentage of hairy roots with prehaustoria (A) and prehaustoria number per plant (B) are shown. Hairy roots with strong GFP signals were selected to minimize chimeric affects. Empty vector transformed WT was used as a control. Data represent mean ± SE from three representative experiments. n =16~30. Student's t test, unpaired, two-tailed (**P < 0.01).

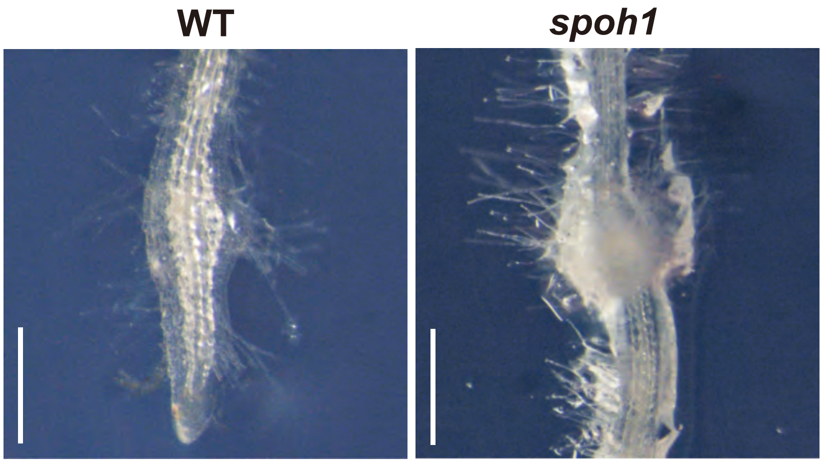


**fig. S16. Prehaustorium formation in WT and *spoh1***
Representative images of a WT root tip treated with DMBQ forming a prehaustorium (left) and a spontaneous prehaustorium in *spoh1* at the maturation zone (right). Scale bar: 500 µm.

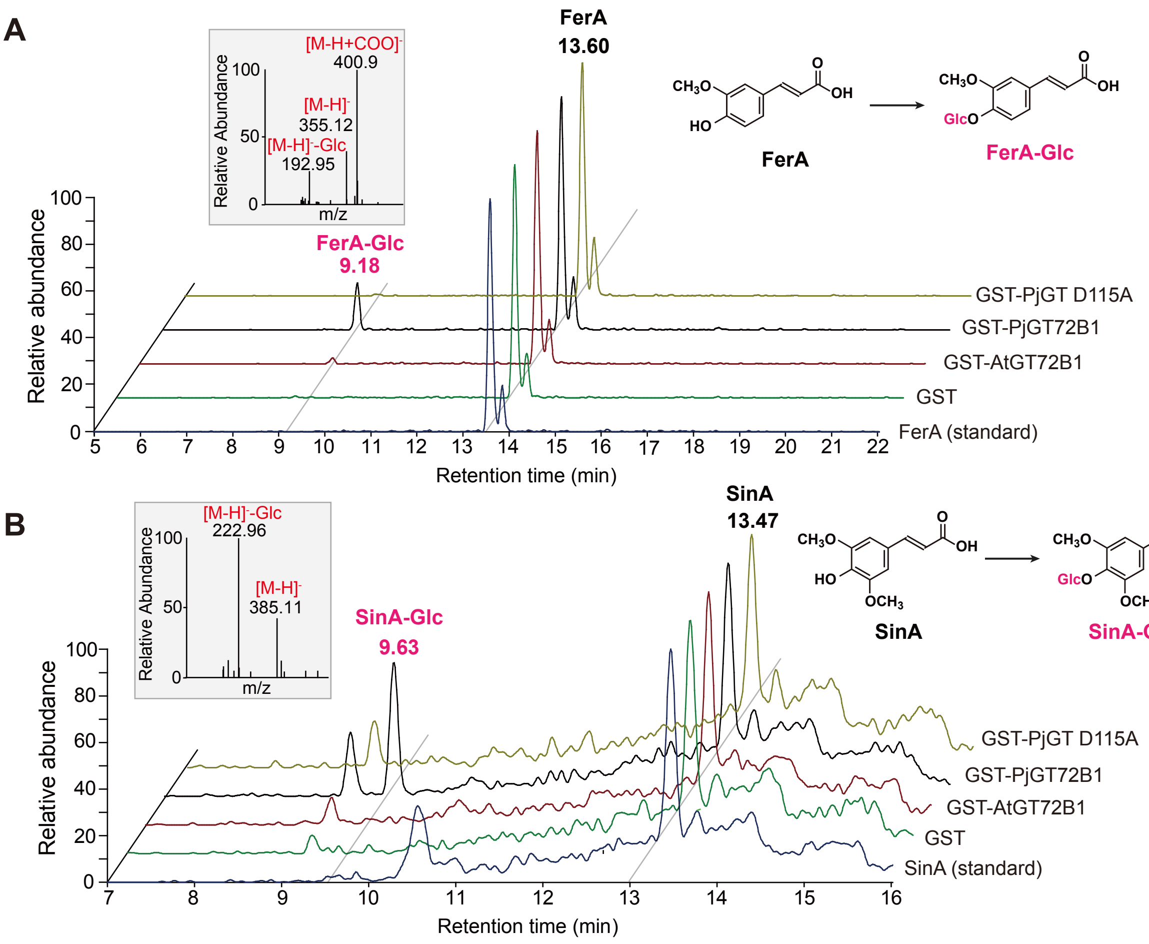


**fig. S17. Prehaustorium formation in WT and *spoh1***

(A, B) LC-MS chromatograms of enzyme reaction products using ferulic acid (A) or sinapic acid (B) as substrates. GST alone and the GST-PjGT72B1 D115A mutant (PjGT D115A) served as negative controls. STD denotes the chromatogram of the standard substrate. Insets display the corresponding MS peaks.

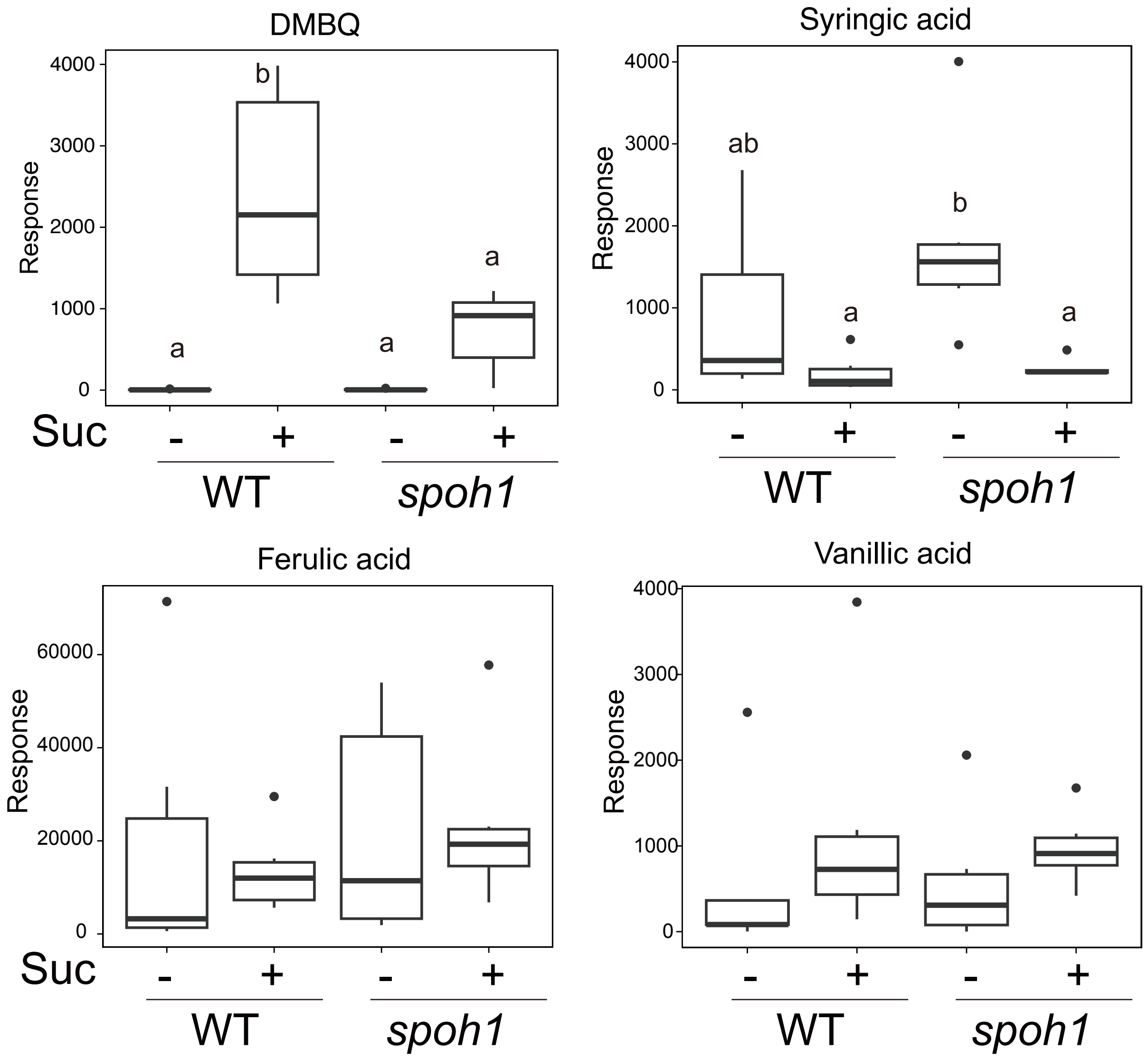


**fig. S18. Quantification of HIFs in root exudates**
The indicated HIFs were analysed by MRM-LC-MS/MS in WT and *spoh1* root exudates incubated with or without 4% sucrose. Response indicates peak area corresponding to the indicated compounds. n=5~6. Different letters indicate significant differences determined by Tuley HSD test ($p<0.05$). The absence of letters in a panel indicates there are no significant differences between the samples.

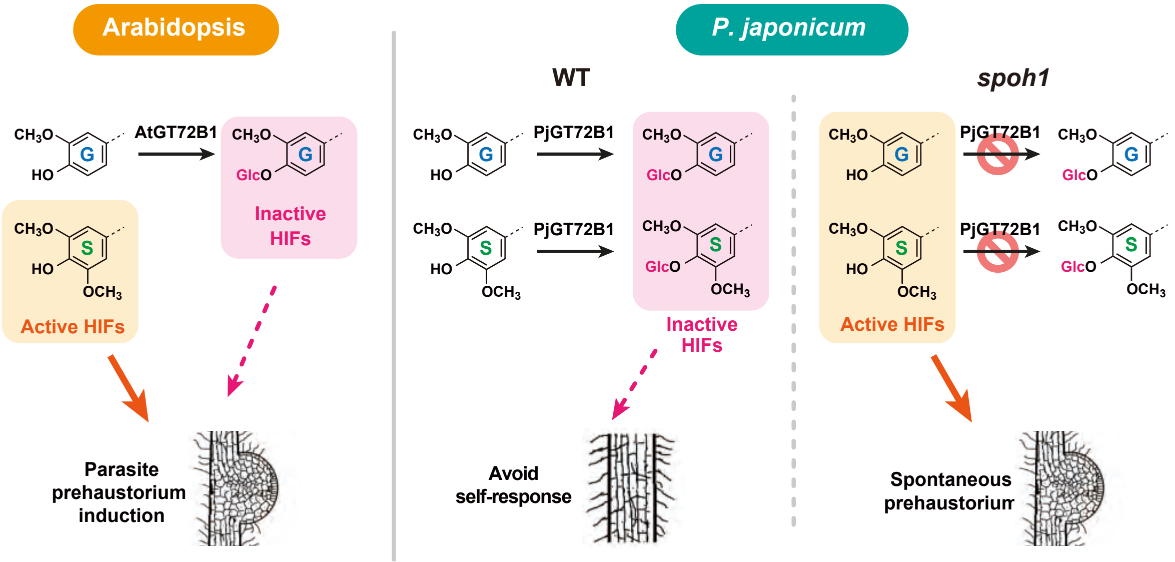


**fig. S19. A model for kin avoidance of parasitic plants through HIF glucosylation**
In Arabidopsis (a host plant), AtGT72B1 glucosylates G-type lignin-related phenolics but not S-type, which therefore remain as active HIFs. In *P. japonicum* WT, both G- and S-type phenolics are glucosylated by PjGT72B1 and therefore no longer active as HIF. In *spoh1*, PjGT72B1 is mutated, and therefore active HIFs are produced by the parasite, which induce spontaneous prehaustorium formation.

**Table S1 Segregation of spontaneous haustorium phenotypes in the F1 and F2 generations of the *spoh1* mutant backcrossed with the wild type**

| Cross (♀ × ♂) | Total No. | Normal | Plant with spontaneous Prehaustoria | Ratio | χ2 | p-value |
|---|---|---|---|---|---|---|
| F1 generation *spoh1* × WT | 56 | 51 | 5 | -- | -- | -- |
| F2 generation *spoh1* × WT | 988 | 765 | 223 | 3:1 | 3.1093 | 0.0778 |

* p>0.05, accept the null hypothesis

**Table S2 Primers used for this study**

| Primer name | Sequence (5'-3') |
|---|---|
| <For Cloning> | |
| pUB-PjGT-F | TTGATGTGATTACAGATGGCGGCCACACCA |
| pUB-PjGT-R | AGTCACTATGGTCGATCAAATGGAAGCACTATTACTCT |
| pUB-WAS-F | TTGATGTGATTACAGATGGCGGCCACACCA |
| pUB-WAS-R | AGTCACTATGGTCGATTAAATCCTCGGGATCGACTCCGAA |
| pUB-ATGT-F | TTGATGTGATTACAGATGGAGGAATCCAAAACACC |
| pUB-ATGT-R | AGTCACTATGGTCGATTAGTGGTTGCCATTTTGC |
| pUB-11317-F | TTGATGTGATTACAGATGGGCCACCTCATACC |
| pUB-11317-R | AGTCACTATGGTCGATCATTCTAGTCCAATCCATT |
| proGT-F | TCGGATCCGGAGCCCCGTGGGCAATGTTCG |
| proGT-R | TTGCTCACCATTCCCGGACGAAGAGGAGAGAGAAAAAGAAGAG G |
| pDE-PjGT-F | ACAAGTTTGTACAAAATGGCGGCCACAC |
| pDE-PjGT-R | AGTCACTATGGTCGATCAAATGGAAGCACTATTACTCT |
| pDE-ATGT-F | ACAAGTTTGTACAAAATGGAGGAATCCAAAACAC |
| pDE-AtGT-R | AGTCACTATGGTCGATTAGTGGTTGCCATTTTG |
| <For CRISPR> | |
| guide-PjGT-1a | CGGACGGCGTCGCATATCGA |
| guide-PjGT-1b | CCCCATGTGCTTCTCTGAAG |
| guide-PjGT-2a | GCGGAGAGGGGCCCGTCGGT |
| guide-PjGT-2b | CCCCTCGGGCAGATAGGCCA |
| guide-11317-1a | GCACACCTTCCACGCGGTTA |
| guide-11317-1b | CGGTATGGGTAATAGCCCGC |
| guide-11317-2a | CAGCGCTCCACGACCTGAAC |
| guide-11317-2b | GCAAGTCAGATCCGCGAACC |
| <For qRT-PCR> | |
| Pj_PTB-F | TCCGATGCAACAAGCTCCTGGG |
| Pj_PTB-R | ATGTGCCAGGAGCGGACACAAA |
| PjGT-QRT-F | ACGTCTCTTTCGGGAGTGGT |
| PjGT-QRT-R | TAGGGCATCTAAGCACCCAC |
| PjWAS-QRT-F | TTGGCGTGCCTTTCTCATTTT |
| PjWAS-QRT-R | GGTGGGAGGAGAGATTACGG |
| AtGT-QRT-F | TGACTCGTTCAAACCCGGAG |
| AtGT-QRT-R | AAAGCGTCCGTACCGAAGAG |

**Data S1. SNPs detection from F2 genome sequencing**

**Data S2. DEGs detected comparison of WT and *spoh1* in water-agar condition**

**Data S3. DEGs detected comparison of WT and *spoh1* in sucrose-agar condition**

**Data S4. Upregulated genes in *spoh1* in both conditions**

**Data S5. Downregulated genes in *spoh1* in both conditions**

**Data S6. SOM clustering analysis**
